# Narcissus reflected: gray and white matter features joint contribution to the default mode network in predicting narcissistic personality traits

**DOI:** 10.1101/2024.01.21.576578

**Authors:** Khanitin Jornkokgoud, Teresa Baggio, Richard Bakiaj, Peera Wongupparaj, Remo Job, Alessandro Grecucci

**Affiliations:** Department of Research and Applied Psychology, Faculty of Education, Burapha University, Thailand; Department of Psychology and Cognitive Sciences (DiPSCo), University of Trento, Italy; Centre for Medical Sciences (CISMed), University of Trento, Italy; Department of Psychology, Faculty of Humanities and Social Sciences, Burapha University, Thailand

**Author notes:** Corresponding author; Department of Psychology and Cognitive Sciences (DiPSCo), University of Trento, Corso Bettini, 31, Rovereto (TN) 38068, Italy. Conflicts of Interest: The authors declare no conflict of interest.

**Keywords:** narcissism, narcissistic personality traits, parallel ICA, Random Forest, machine learning

## Abstract

Despite the clinical significance of narcissistic personality, its neural bases have not been clear yet, primarily due to methodological limitations of the previous studies, such as the low sample size, the use of univariate techniques and the focus on only one brain modality. In this study, we employed for the first time a combination of unsupervised and supervised machine learning methods, to identify the joint contributions of gray (GM) and white matter (WM) to narcissistic personality traits (NPT). After preprocessing, the brain scans of 135 participants were decomposed into eight independent networks of covarying GM and WM via Parallel ICA. Subsequently, stepwise regression and Random Forest were used to predict NPT. We hypothesize that a fronto-temporo parietal network mainly related to the Default Mode Network, may be involved in NPT and white matter regions related to these regions. Results demonstrated a distributed network that included GM alterations in fronto-temporal regions, the insula, and the cingulate cortex, along with WM alterations in cerebellar and thalamic regions. To assess the specificity of our findings, we also examined whether the brain network predicting narcissism could predict other personality traits (i.e., Histrionic, Paranoid, and Avoidant personalities). Notably, this network did not predict these personality traits. Additionally, a supervised machine learning model (Random Forest) was used to extract a predictive model to generalize to new cases. Results confirmed that the same network could predict new cases. These findings hold promise for advancing our understanding of personality traits and potentially uncovering brain biomarkers associated with narcissism.

## 1. Introduction

The term narcissism encompasses a spectrum of traits ranging from healthy personality features to pathological states, potentially culminating in Narcissistic Personality Disorder (NPD) (Maj, 2005). High levels of narcissism could significantly impact mental health and interpersonal relationships(George & Short, 2018). In the general population, narcissism prevalence estimates range from 0% to 5.3% (Ronningstam, 2009) and 18.1% (Pourramzani & Monajemi, 2021), while NPD is believed to affect approximately 1% to 15% of the US population in clinical settings (Mitra & Fluyau, 2020). Diagnosing NPD can be challenging, particularly when it coexists with other mental disorders (Mitra & Fluyau, 2020). As a matter of fact, patients with NPD may exhibit symptoms shared with other personality disorders, including histrionic (for their need to be the focus of others’ attention), paranoid (when attacking others to maintain their self-esteem), and avoidant traits (avoiding certain situations to protect the fragile self) (American Psychiatric Association, 2013; McGonigal & Dixon-Gordon, 2020; Ponzoni et al., 2021).

Both clinical and non-clinical investigations have explored narcissistic traits (Nenadić et al., 2021), primarily relying on validated questionnaires, such as the Narcissistic Personality Inventory (NPI) (Raskin & Hall, 1979). However, research into the structural and functional correlates of narcissistic traits remains limited, with most studies having used separate analyses of a single neuroimaging modality, specifically gray matter (GM) or white matter (WM) (Chester et al., 2016; Jornkokgoud et al., 2023; Nenadić et al., 2021), and having employed univariate methods.

Despite the limited investigations conducted so far, narcissistic personality traits (NPT) have been associated with structural and functional alterations in specific brain regions, including the prefrontal cortex, the dorsal anterior cingulate and subgenual cingulate cortices, and the insula (Nenadić et al., 2021). For instance, while Schulze et al. (2013) reported a reduced gray matter volume in fronto-paralimbic brain regions in NPD patients, such as the rostral and median cingulate cortex, dorsolateral and medial parts of the prefrontal cortex, another study by Yang et al. (2015) found a positive correlation between female NPI scores and regional GM volume in the right superior parietal lobule. Mao et al. (2016) utilized a vertex-wise general linear model (GLM) to investigate the relationship between scores on the pathological narcissism inventory (PNI), cortical thickness (CT), and cortical volume (CV). These findings revealed that individual differences in pathological narcissism are linked to specific gray matter features, encompassing multiple regions within the social brain network and the right dorsolateral prefrontal cortex, a part of the central executive network. However, the studies mentioned above may suffer from major limitations, including a low number of participants (i.e., N=34) and reliance on massive univariate statistical approaches that do not consider the joint contribution of multiple voxels, and thus failing to characterize the effect of interest from a network perspective.

In a recent effort to overcome some of these limitations, Jornkokgoud and colleagues (2023) employed a multivariate supervised machine learning approach, specifically Support Vector Machine in the form of Kernel Ridge Regression. Their study successfully identified a GM network including the middle frontal gyrus, angular gyrus, Rolandic operculum, Rectus, and Heschl’s gyrus as a predictive indicator of NPT. In the similar vein, using resting-state properties of the brain, a medial prefrontal-frontal network was found to be a predictor of narcissism, including areas such as the medial prefrontal cortex (mPFC), lateral prefrontal cortex (PFC), amygdala, and the dorsal anterior cingulate cortex (dACC) (Feng et al., 2018).

For what concerns the white matter contribution to narcissistic personality, few studies have examined fiber tract integrity using Diffusion-Tensor Imaging (DTI), a Magnetic Resonance Imaging (MRI) technique that quantifies diffusion properties in white matter (Assaf & Pasternak, 2008). Nenadić and colleagues (2015) reported decreased fractional anisotropy (FA, an index of white matter integrity) in the right frontal and temporal lobe, right anterior thalamic radiation and the right brain stem. Chester and colleagues (2016) also observed an association between narcissism and reduced WM integrity in fronto-striatal pathways. Although DTI is the most commonly used technique to assess WM fibers integrity (Assaf & Pasternak, 2008; De Erausquin & Alba-Ferrara, 2013; Radwan et al., 2022), it is sensitive to noise and, because of this, region-of-interest-driven protocols are often poorly reproducible (Assaf & Pasternak, 2008; Baggio et al., 2023). Moreover, DTI assumes a Gaussian diffusion in white matter and employes a single diffusion tensor per pixel, limiting the analysis to specific tracts. (Assaf & Pasternak, 2008). Recent evidence suggests that raw white matter concentration can successfully be used to characterize WM abnormalities, allowing for the evaluation of widely distributed alterations not only confined to specific tracts (Baggio et al., 2023). Moreover, by using multivariate methods instead of univariate methods, WM can be characterized in its network-like organization (O’Muircheartaigh & Jbabdi, 2018; Peer et al., 2017).

Most research has focused on gray or white matter networks separately, with the joint contribution of GM and WM to narcissistic traits remaining unexplored. Given that GM and WM possibly share similar genetic influences (Baggio et al., 2023; Panizzon et al., 2012) and the integrity of one modality affects the other (Jacków-Nowicka et al., 2021), it is essential to investigate them together in order to identify potential joint contributions to narcissistic personality.

Accordingly, the current investigation aimed to characterize the joint GM and WM contributions to narcissistic personality traits, to assess their specificity in predicting narcissistic traits versus other personality traits, and build a predictive model to predict new individuals. Machine learning (ML) approaches have gained increasing public and academic traction in neuroimaging research, showcasing promising outcomes in predicting cognition and behavioral patterns in mental disorders (Caria & Grecucci, 2023; Ghomroudi et al., 2023; Grecucci, Rastelli, et al., 2023; Grecucci, Sorella, et al., 2023). Among the diverse ML techniques available, Independent Component Analysis (ICA) stands out as a valuable tool for unsupervised blind source separation (Douglas et al., 2013). The ICA demonstrates its ability to unveil distinct neural circuits throughout the entire brain by leveraging structural MRI data from individual subjects (Ghomroudi et al., 2023; Lapomarda et al., 2021; Sorella et al., 2019). Unsupervised ML methods excel in revealing concealed structures within image collections or identifying sub-populations in extensive cohorts. In contrast, supervised ML typically seeks to establish connections between brain images and behavioral or clinical observations, often employing decoding or encoding approaches (Abraham et al., 2014). Unsupervised ML, also known as ‘data-driven,’ operates solely on input data without requiring the presence of predefined target variables, distinguishing it from supervised learning (Claude et al., 2020). These approaches have the potential to generate large-scale hypotheses regarding GM-WM networks, elucidate brain-behavior correlations, and uncover underlying structures and mechanisms (Vu et al., 2018).

Parallel Independent Component Analysis (p-ICA), an unsupervised method based on Independent Component Analysis (ICA), facilitates the integration of different modalities, such as gray and white matter, by uncovering maximally independent components for each imaging modality and establishing their interrelationships within a single algorithm (Grecucci, Dadomo, et al., 2023; Kubera et al., 2019; Liu et al., 2008; Yang et al., 2019). Researchers have developed multivariate data fusion techniques aimed at extracting comprehensive and complementary information from multimodal neuroimaging data. For instance, p-ICA was applied to investigate co-varying patterns of gray matter (GM) and white matter (WM) tissue damage in patients with early relapsing-remitting multiple sclerosis, shedding light on its implications for the progression of chronic neuroinflammatory diseases (Muthuraman et al., 2020). More recently, Baggio and colleagues (2023) leveraged p-ICA to explore the co-varying patterns of GM and WM in trait anxiety, while Grecucci and colleagues (2023) employed the same technique to identify brain abnormalities in borderline personality disorder. Consequently, this approach proves to be highly effective for conducting a sophisticated analysis of structural networks associated with specific psychological variables. However, one limitation of unsupervised ML methods such as Parallel ICA is that it cannot be used to predict new cases. Supervised ML methods, are instead the best option for this task. In the present study we used a supervised ML method known as Random Forest to build a predictive model based on the output of the Parallel ICA results. Recent studies in neuroimaging have made use of statistical regression methods, such as stepwise regression to build predict psychological variables from neural features (Radetz et al., 2020; Schmidt et al., 2023). However, regression can only draw inferences about the population considered and does not return a measure of generalizability to new unobserved cases. Supervised ML methods such as Random Forest regression are by contrast excellent ways to extract a predictive model and allow for generalization of the results outside the sample considered via the cross validation procedure (Grecucci, Dadomo, et al., 2023; Jollans et al., 2019). In the present study we intend to use both approaches (stepwise regression and Random Forest) to understand the brain bases of NPT.

Building upon our previous study, the current study has three main objectives. The primary objective of this study was to discover joint GM and WM contributions to unveil the intricate structural networks associated with NPT. In addition, we aimed to validate the specificity of our findings by assessing the possibility to predict other personality traits that may share some common features with narcissistic personalities, such as the histrionic (Cluster B), the avoidant (Cluster C), and the paranoid personalities (Cluster A). We expected to find a co-varying GM-WM network able to predict the severity of NPT but not of other personality traits. This network should encompass GM regions in frontal and temporal areas, including the middle frontal gyrus, angular gyrus, Rolandic operculum, Rectus, and Heschl’s gyrus, as well as the cingulate and insula. Of note these regions should be ascribed to the Default Mode network. The default mode network (DMN) is activated during rest and not only consisted of two major hubs, corresponding to the medial prefrontal cortex and the posterior cingulate cortex (PCC), but also temporo-pariatel regions (Langerbeck et al., 2023). It also includes the hippocampus and the cingulate gyrus, which are known to play a role in semantic memories, internal thoughts, self-monitoring and emotion regulation (Menon, 2011). One intriguing hypothesis is that abnormalities inside the regions ascribed to the DMN may be related to NPT as well. Furthermore, we expected to identify WM abnormalities in fronto-temporal regions, along with thalamic and cerebellar areas, consistent with prior research on personality (Assaf & Pasternak, 2008; Chester et al., 2016), and known to support the DMN functionality (Teipel et al., 2010). These regions may subserve diverse functions related to personality, from fronto-temporal distorted representations of self and others (Hu et al., 2016), to the cingulate and insular regions contributing to unusual emotional experiences (Medford & Critchley, 2010), to the cerebellum’s role in various forms of psychopathology (Phillips et al., 2015). Although we aimed to use structural data, several previous studies have shown that resting state macro-networks are also present at a structural level when data are analyzed with ICA-based methods (Baggio et al., 2023; Grecucci et al., 2022; Meier et al., 2016; Vanasse et al., 2021). In other words, the well-known macro-networks are present at both functional and structural levels (Vanasse et al., 2021).

Last but not least, the third objective of this study was to extract a predictive model to generalize to new cases via Random Forest regression. We expect to find that the same network found in the stepwise regression, should be the main predictor of new cases in the Random Forest analysis.

## 2. Methods

### 2.1. Participants

For this study, we utilized data from the MPI-Leipzig Mind Brain-Body dataset, which is accessible through the OpenNeuro database (Accession Number: ds000221) (Babayan et al., 2020). This dataset encompasses MRI and behavioral data gathered from 318 participants who participated in the project conducted by the Max Planck Institute (MPI) of Human Cognitive and Brain Sciences in Leipzig. The study was carried out under the authorization of the ethics committee at the University of Leipzig (Protocol ID: 154/13-ff) (Babayan et al., 2019).

For the purposes of this study, we meticulously selected data from a subset of 135 healthy participants (64 females and 71 males, Mean age range = 31.94, SD = ± 15.06) who participated in the Neuroanatomy & Connectivity Protocol. Eligibility for inclusion in our study was contingent upon the participants’ good health, absence of any medication use, and no history of substance abuse or neurological diseases.

### 2.2. Personality Styles and Disorders Inventory (PSDI)

In this study, we used the PSDI (German version: Persönlichkeits-Stil-und Störungs-Inventar, PSSI), taking into consideration only the following subscales: narcissist (Mean score = 12.14, SD = ± 4.76), histrionic (Mean score = 12.92, SD = ± 4.40), insecure/avoidant (Mean score = 11.50, SD = ± 4.37) and paranoid (Mean score = 10.04, SD = ± 4.34). The PSDI is a validated self-report inventory that measures various personality styles and can provide insights into potential personality disorders, particularly when extreme scores are observed. Developed and revised by Kuhl and Kazén (2009), the inventory consists of 140 items organized into 14 subscales. In our study, we considered the scores of the ‘narcissist’ subscale of the PSDI to examine the hypothesis that individual differences in narcissistic personality traits can be predicted from brain circuits. The PSDI shows a theoretically extraordinarily coherent network of relationships with a large number of clinical and non-clinical behavioral characteristics (e.g., suicidality, depression, psychosomatic symptoms, the Big Five personality traits, and the sixteen personality factors). This extensive network demonstrates a good construct validity of the inventory. Furthermore, the PSDI employs objective procedures and analyses, and its consistency coefficients (Cronbach’s alpha) range between α = .73 and .85. Test-retest coefficients vary between *r* = .68 and .83, indicating good test-retest reliability. The construct validity of the inventory is considered acceptable for both clinical and non-clinical behavior (Kuhl & Kazén, 2009).

### 2.3 MRI Data Acquisition

The MPI-Leipzig Mind Brain-Body dataset comprises quantitative T1-weighted, functional, resting state, and diffusion-weighted images acquired at the Day Clinic for Cognitive Neurology of the University Clinic Leipzig and the Max Planck Institute for Human and Cognitive and Brain Sciences (MPI CBS) in Leipzig, Germany (Babayan et al., 2019). For the purpose of our research, we only considered the T1-weighted images. Magnetic Resonance Imaging (MRI) was performed on a 3T Siemens MAGNETOM Verio scanner (Siemens Healthcare GmbH, Erlangen, Germany) with a 32-channel head coil. The MP2RAGE sequence consisted of the following parameters: sagittal acquisition orientation, one 3D volume with 176 slices, TR = 5000 ms, TE = 2.92 ms, TI1 = 700 ms, TI2 = 2500 ms, FA1 = 4°, FA2 = 5°, pre-scan normalization, echo spacing = 6.9 ms, bandwidth = 240 Hz/pixel, FOV = 256 mm, voxel size = 1 mm isotropic, GRAPPA acceleration factor 3, slice order = interleaved, duration = 8 min 22 s.

### 2.4 MRI Data Preprocessing

We performed the MRI preprocessing on all anatomical images through SPM12 and the Computational Anatomy Toolbox (CAT12), in the environment of MATLAB. After the manual re-orientation through the anterior commissure, images were segmented into gray matter, white matter, and cerebrospinal fluid. In this study, only gray and white matter were further used for the successive steps. A diffeomorphic anatomical registration exponential Lie algebra (DARTEL) approach was then applied to normalize each subject’s gray matter image to the average DARTEL template and to the Montreal Neurological Institute (MNI) space. Finally, a smoothing of 10 was used (Monté-Rubio et al., 2018).

### 2.5 Data fusion unsupervised machine learning (p-ICA)

A parallel independent component analysis (P-ICA) (Liu, Kiehl, et al., 2009; Liu, Pearlson, et al., 2009) was applied to gray and white matter data using the Fusion ICA Toolbox (FIT; http://mialab.mrn.org/software/fit) in MATLAB Version 9.4 (R2018a). Parallel ICA enables to jointly analyze multiple modalities, offering insights into their interplay and interactions (Liu et al., 2008) (see Figure 1). To identify the relationship between gray and white matter modalities, p-values were computed using standard general linear model approaches or bootstrapping for testing the phenotypes against the subject’s loading parameters output of p-ICA. The significance of the correlation coefficients was corrected for all possible combination of pairs between the two modalities (Pearlson et al., 2015). To ensure the reliability of our findings and mitigate the risk of false discoveries due to overfitting, we employed a leave-one-out assessment (Liu, Pearlson, et al., 2009). Thus, given the relatively small number of participants, we conducted 135 runs of parallel ICA with identical parameter settings, each run excluding one subject. This approach allowed us to assess consistency by examining the results from the 135 repetitions.

**Figure 1.**
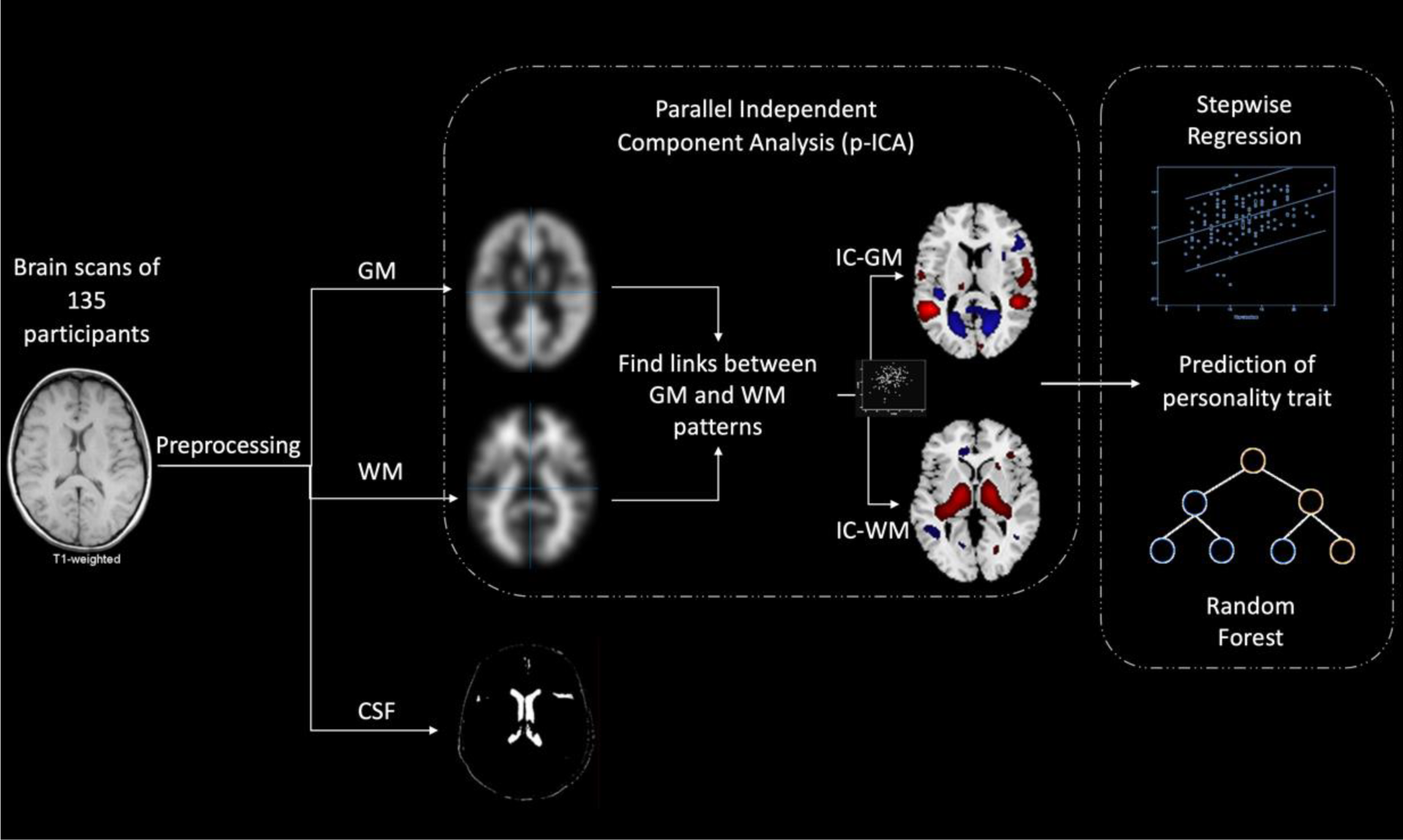
Method. After preprocessing, GM and WM images were entered a data fusion unsupervised machine learning approach (p-ICA), to decompose the brain into joint GM-WM independent networks. Then stepwise regression and Random Forest regression were used to predict narcissistic personality traits.

### 2.6 Characterization of narcissistic personality traits

In our pursuit of understanding the specific brain networks linked to different personality traits, we employed the neural circuits (IC-GMs) uncovered in the previous step in order to predict four distinct personality traits: paranoid (Cluster A), narcissistic and histrionic personality disorders (Cluster B), and avoidant traits (Cluster B). respectively. To delve into this analysis, we harnessed a multiple regression analysis with a stepwise method to examine the loading coefficients of each network for every participant. This analytical approach involved the integration of regressors for both the predictor variable (loading coefficients) and the response variable (four scales extracted from the Personality Styles and Disorder Inventory - PSDI questionnaire). Employing the stepwise regression method allowed us to identify significant predictors and discern their relationships with the targeted personality traits. This data analysis strategy facilitated the prediction of numeric ranges, as it utilized a regression equation incorporating both predictor and response variables. Consequently, in our study, we utilized the loading coefficients as predictors and the four scales from the PSDI questionnaire as responses in the regression analysis.

### 2.7 Predictive model of narcissistic personality traits

We then utilized a supervised machine learning (SML) technique known as Random Forest regression for our analysis. Random Forest is an ensemble learning method designed for regression problems, and it is based on a forest of multiple decision trees (Breiman, 2001; Ho, 1998). The name ‘random forest’ is derived from decision trees, but it distinguishes itself by employing multiple trees and then averaging their performance, a technique known as ‘bagging.’ Notably, random decision forests have proven to be more effective in mitigating overfitting issues compared to many other supervised algorithms (Hastie et al., 2001). To predict personality traits, we used the loading coefficients from each independent network derived through p-ICA as input vectors within the decision trees. These trees made predictions, and the tree with the lowest error rate was chosen as the optimal model. To prevent redundancy and potential collinearity issues that could affect model quality, we exclusively included GM loading coefficients. As a matter of fact, GM and WM are highly correlated, and this issue may cause redundant and unnecessary renders. We employed the ‘hold-out’ method to partition the sample into three subsets: training (70%), validation (15%), and test (15%). The model underwent a structured process, first being trained, followed by validation, and ultimately tested. The statistical results pertain to the model’s performance in predicting new, unobserved cases, providing a measure of its generalization ability as shown in Figure 2.

**Figure 2.**
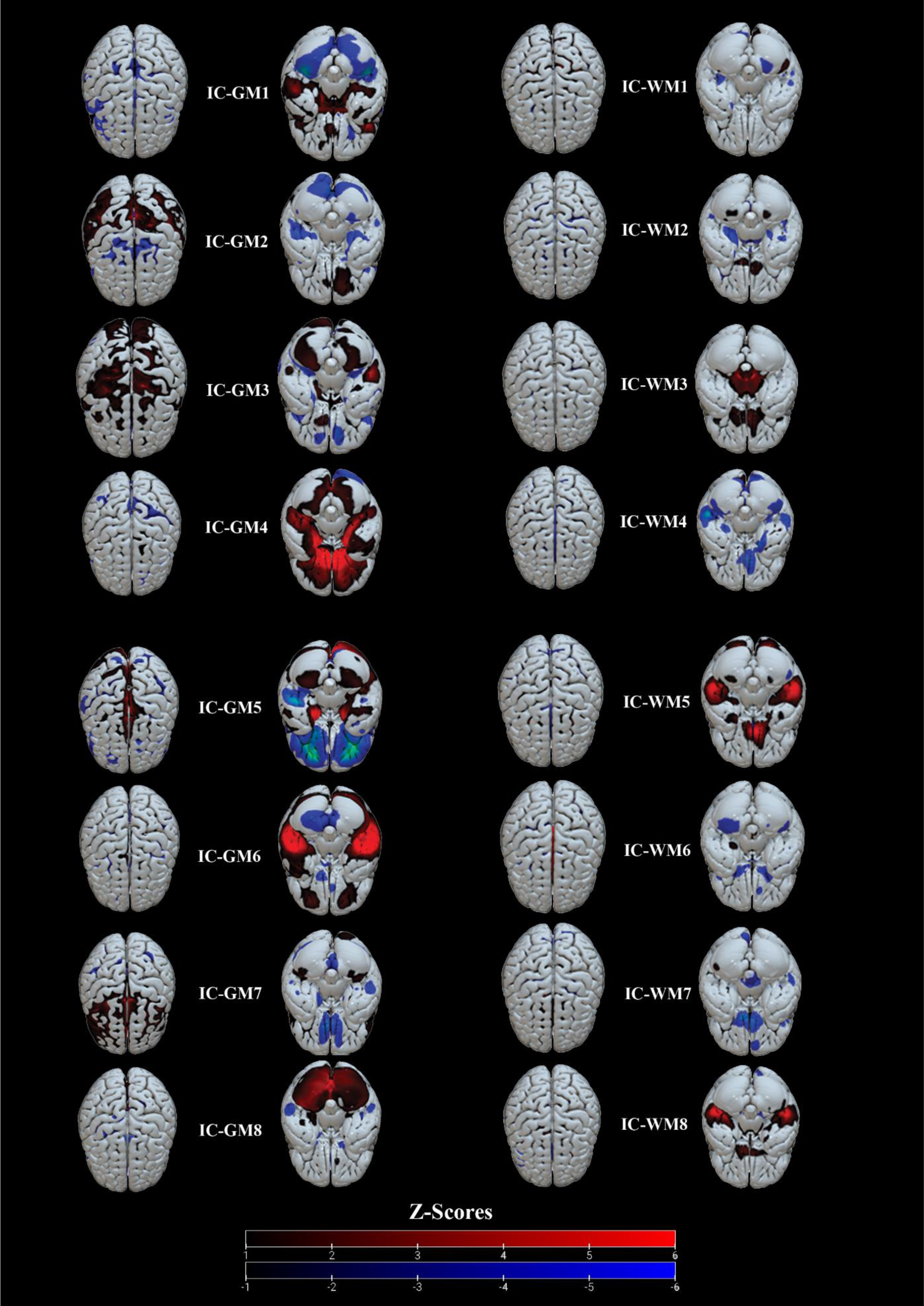
Data Fusion ML results. The eight gray matter-independent components (IC-GM) and the eight white matter-independent components (IC-WM) are shown on the left and right, respectively. Red colors represent regions with positive values, while blue colors represent regions with negative values. The color intensity is computed through z-scores (see the color bar). Each independent component is illustrated from both the superior (left) and inferior (right) positions, and these are displayed on the MNI-ICBM152 atlas using the visualization software Surf Ice (https://www.nitrc.org/projects/surfice/).

## 3. Results

### 3.1. Network decomposition

The Information-theoretic criteria estimated eight independent covarying gray (IC-GM) and eight white (IC-WM) matter networks (see Figure 2). The positive values of these networks indicated increased gray/white matter concentration, whereas negative values indicated decreased concentration. The meaning of the covariation between a gray and white matter component refers to a similar pattern of gray/white matter concentration (i.e., the more gray matter in the circuit IC-GMX, the more white matter in the circuit IC-WMY). The correlation coefficients (*r*) between the independent components of gray matter (IC-GMs) and white matter (IC-WMs) are listed in Table 1.

**Table 1.**
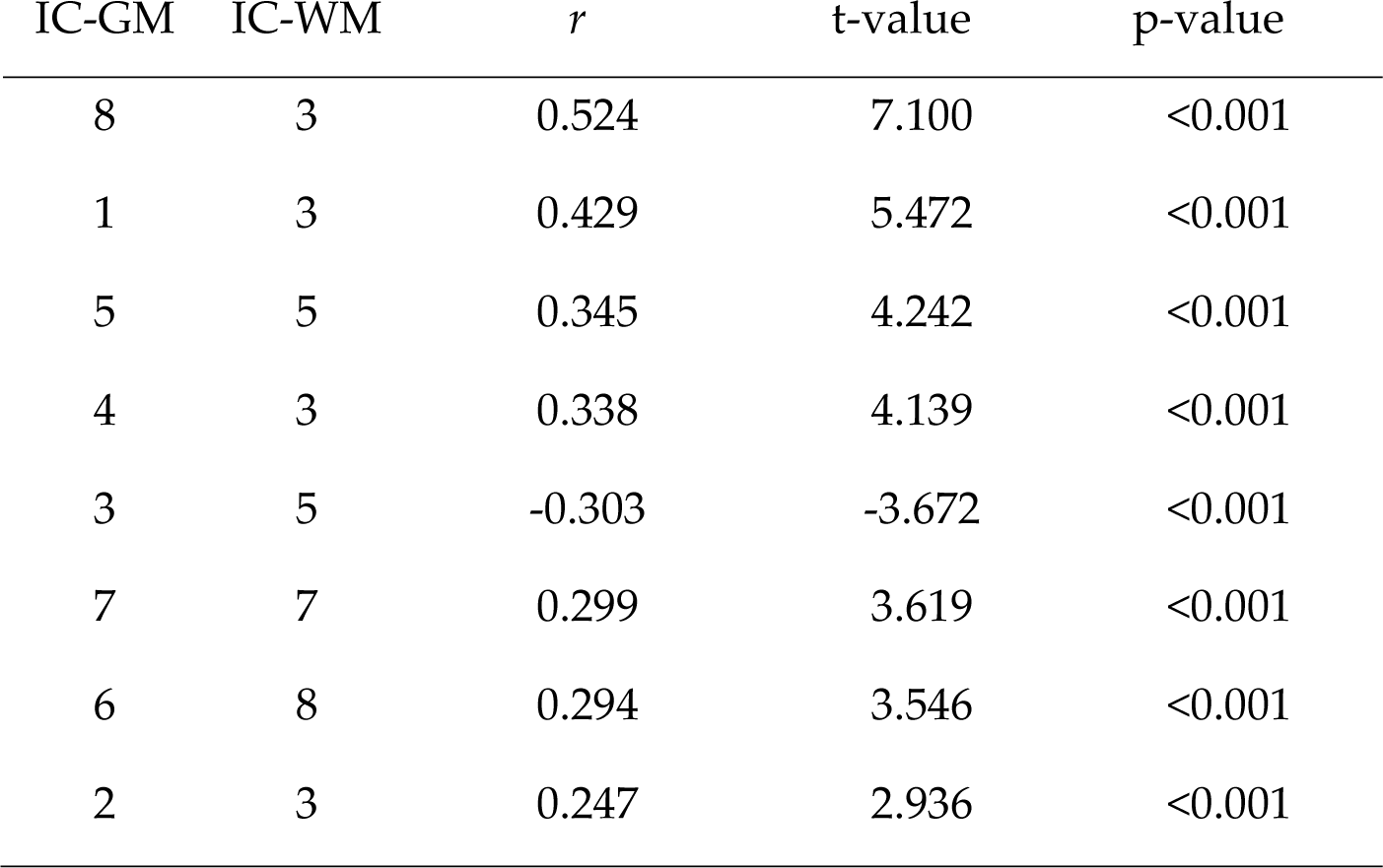
Correlation coefficients between the linked components.

### 3.2 Stepwise regression on loading coefficients

A significant model was found for narcissistic personality traits (*r* = .446, R^2^ = .199, RMSE = 4.279, *p* < 0.001), where only IC-GM2 (*p* < .001) emerged as a surviving predictor, along with IC-WM3. Of note, IC-GM2 and IC-WM3 were substantially correlated (*p*<.001), confirming a joint GM-WM contribution to narcissistic personality traits. Conversely, no significant results were found for histrionic personality traits (*r* =.151, R^2^ =.023, RMSE = 4.368, *p* = .081). Significant results were, however, obtained for paranoid personality traits (*r*=.277, R^2^=.077, RMSE=4.200, *p* = .005) and avoidant personality traits (*r* = .333, R^2^ = .111, RMSE = 4.155, *p* < .001), but from other GM-WM networks: IC-GM3 (*p* = .002) and IC-GM5 (*p* = .041) for paranoid personality traits, and IC-GM1 (*p* = .025) and IC-GM5 (*p* = .007), for avoidant personality traits. This finding confirms the specificity of the network predicting narcissistic traits. Table 2 and Figure 3 report, respectively, the visual representation and the Talairach coordinates of the significant components of narcissistic traits (see supplementary material for the other networks, tables, and plots).

**Figure 3.**
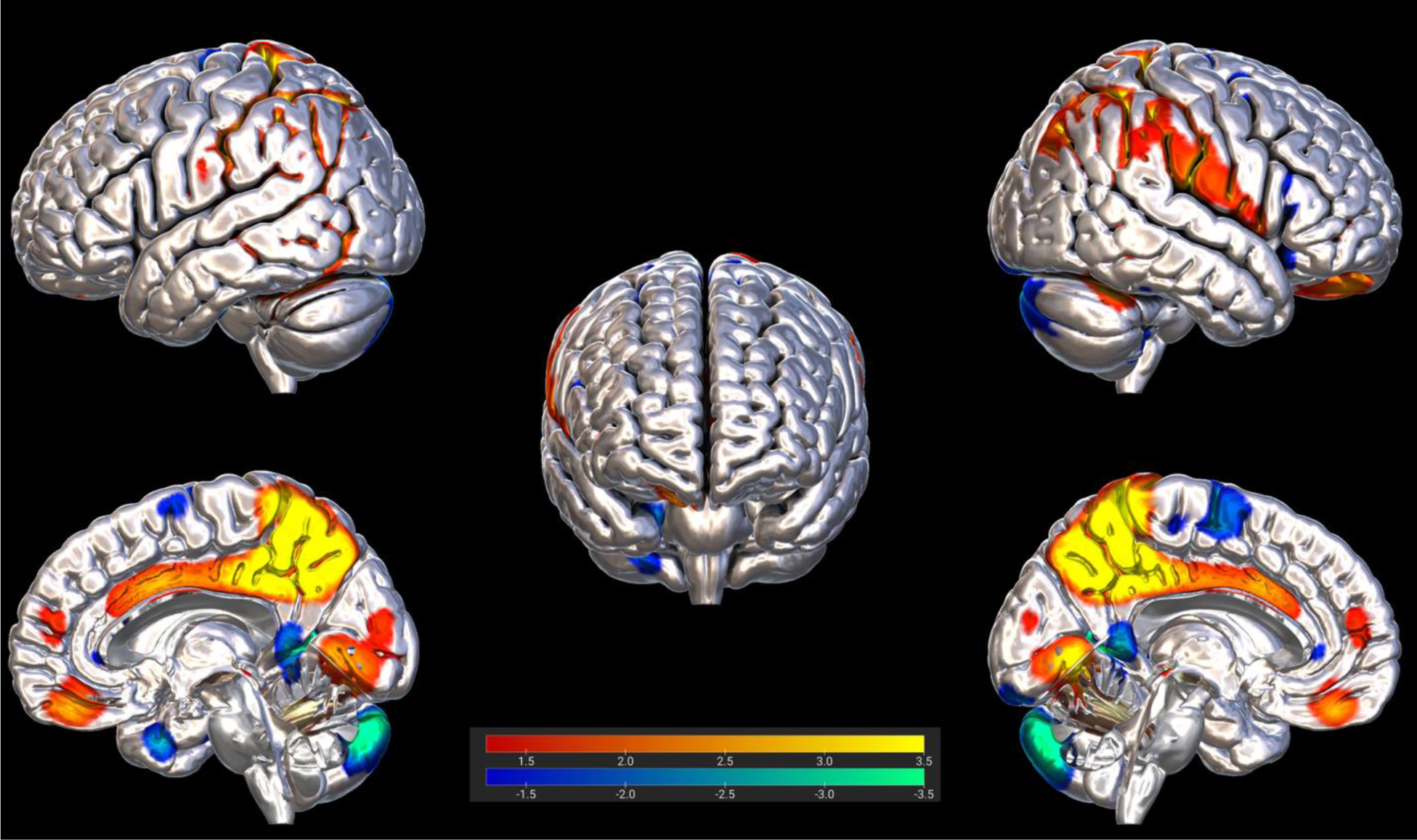
Brain plot of IC-GM2. Significant regions are displayed on the MNI-ICBM152 atlas using the visualization software Surf Ice (https://www.nitrc.org/projects/surfice/). Red and blue colors indicate increased or decreased gray/white matter concentration. The respective color intensity is computed through z-scores (see the color bar).

**Figure 4.**
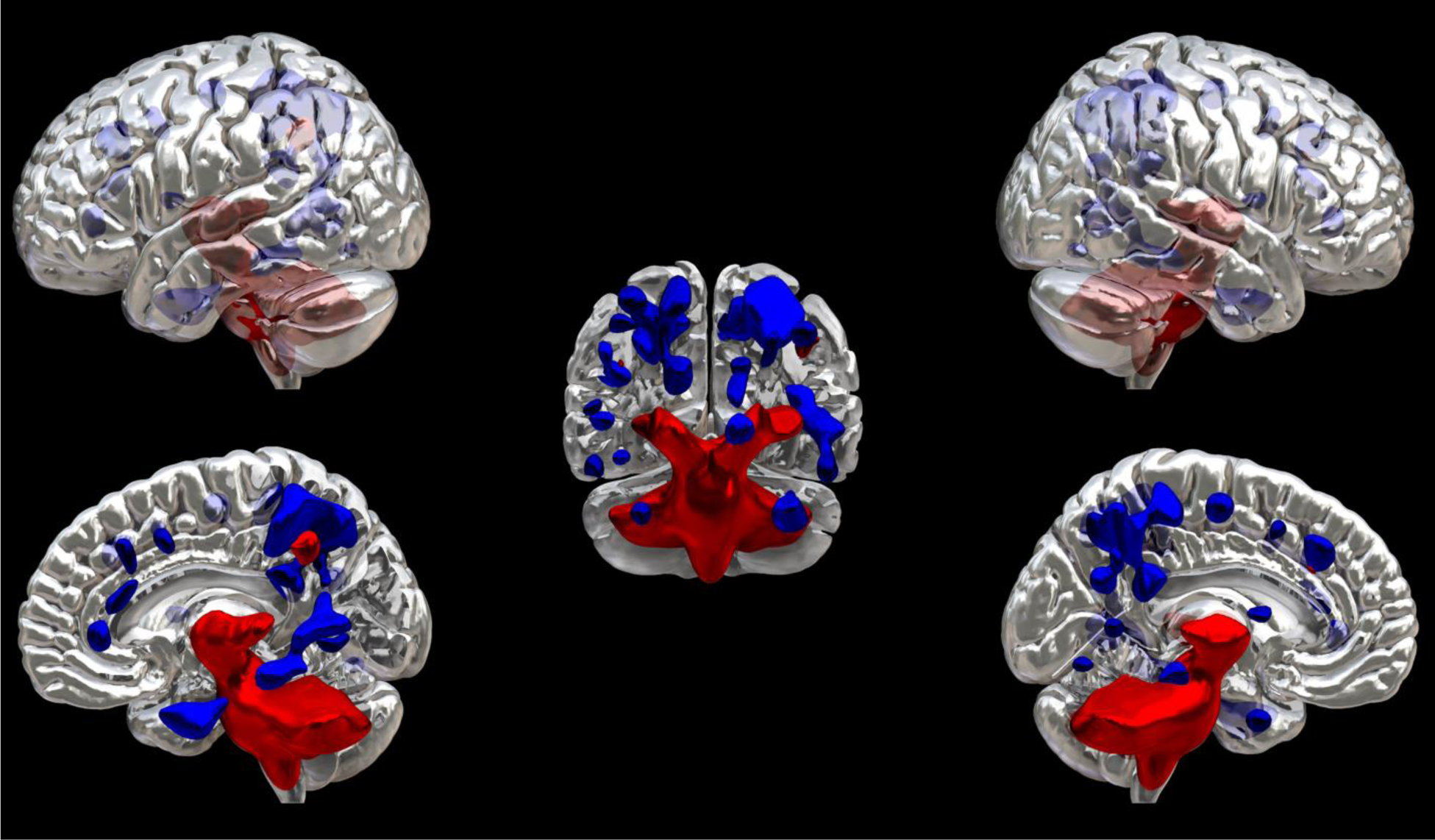
3D mesh plot of IC-WM3. 3D reconstruction in a semi-transparent brain. Significant regions are displayed on the MNI-ICBM152 atlas using the visualization software Surf Ice (https://www.nitrc.org/projects/surfice/). Red and blue colors indicate increased or decreased gray/white matter concentration.

**Table 2.**
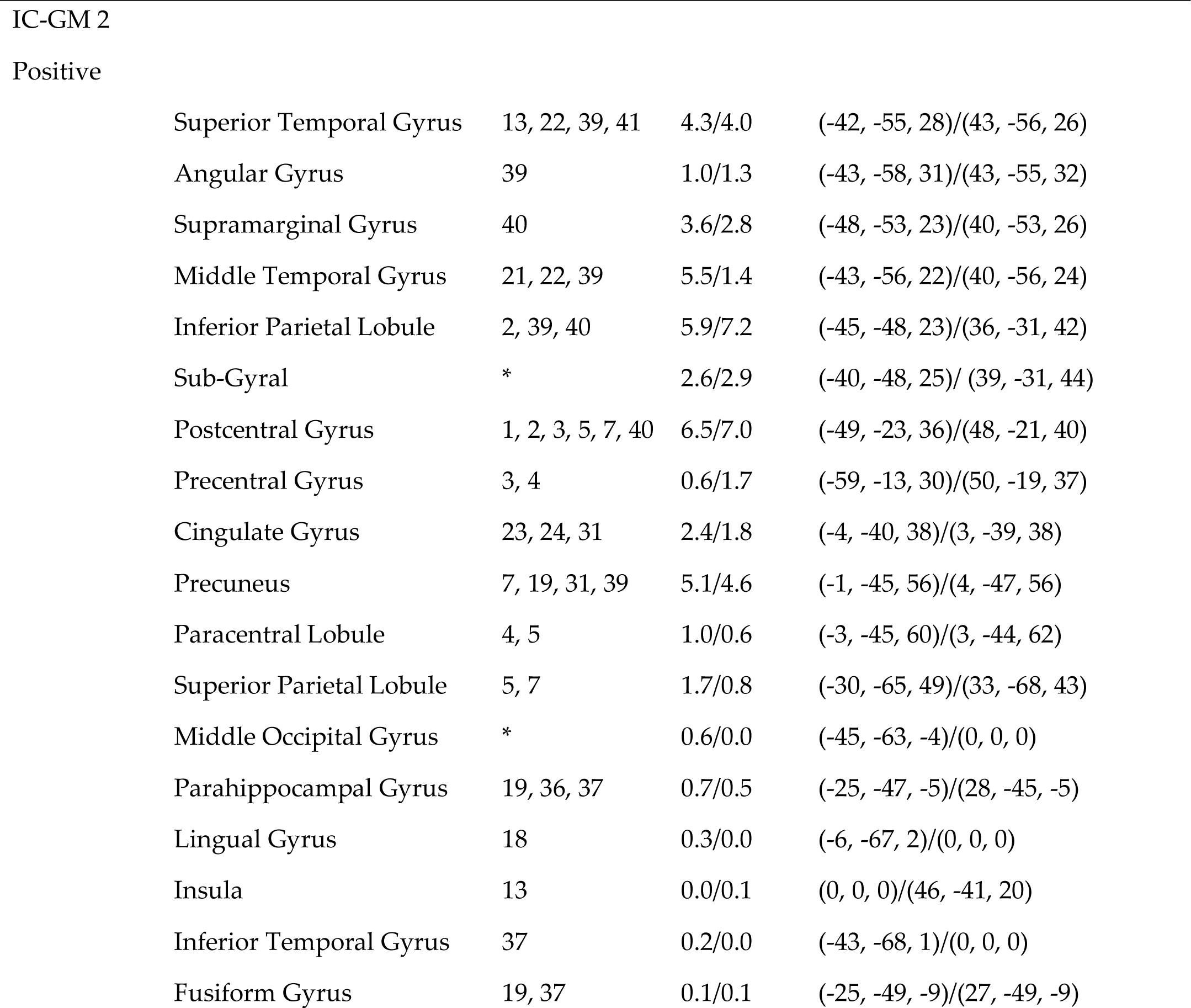
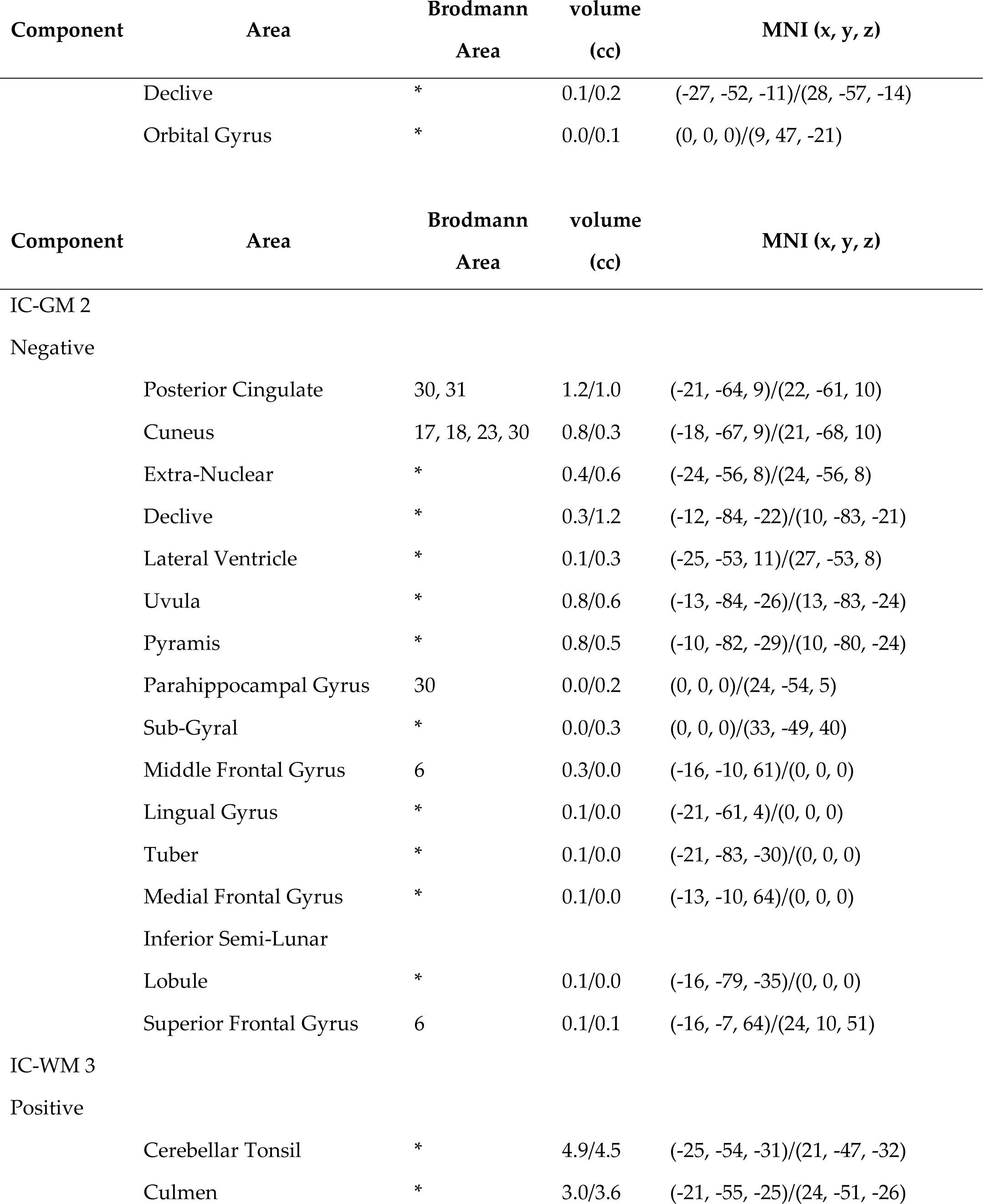
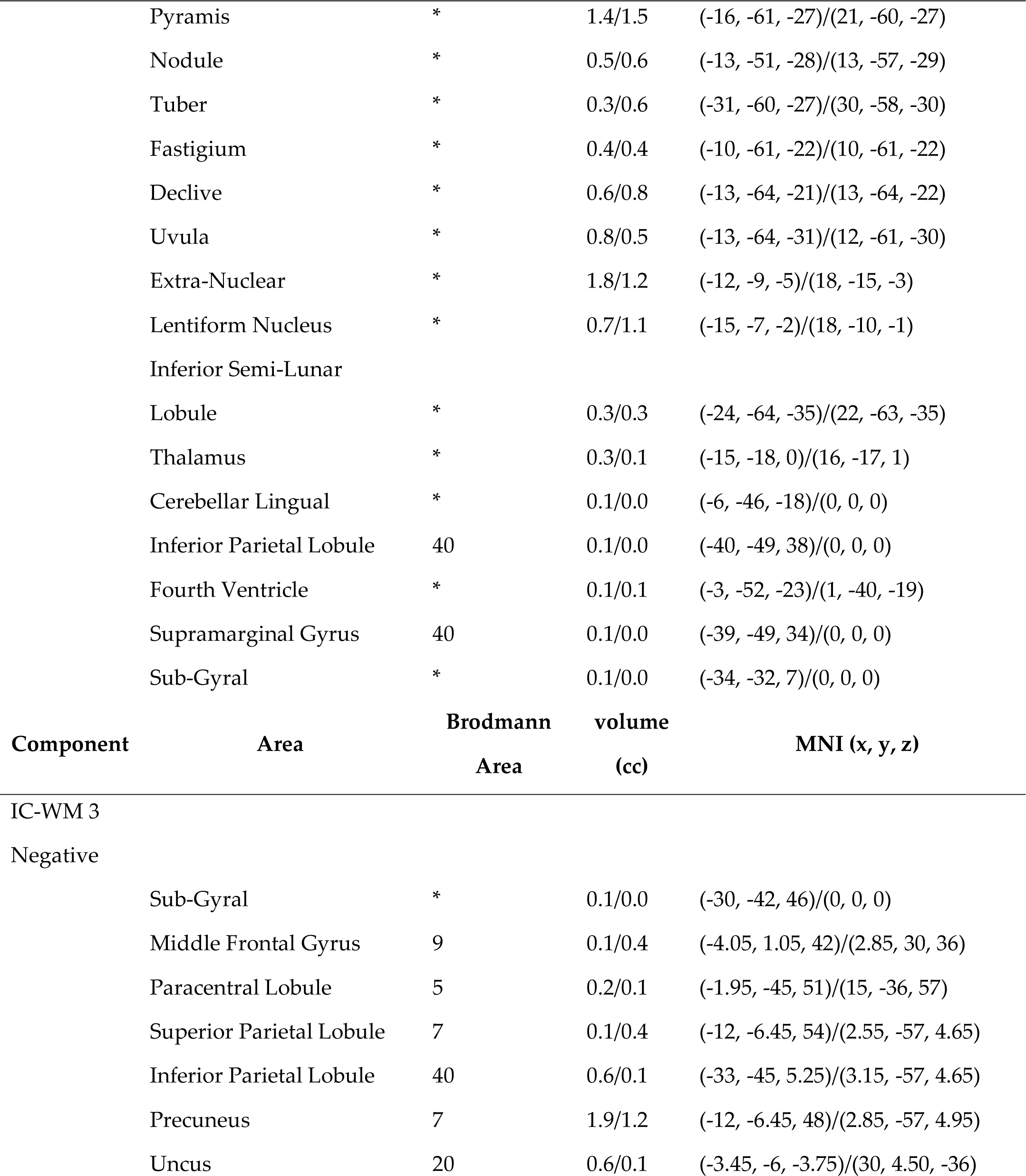
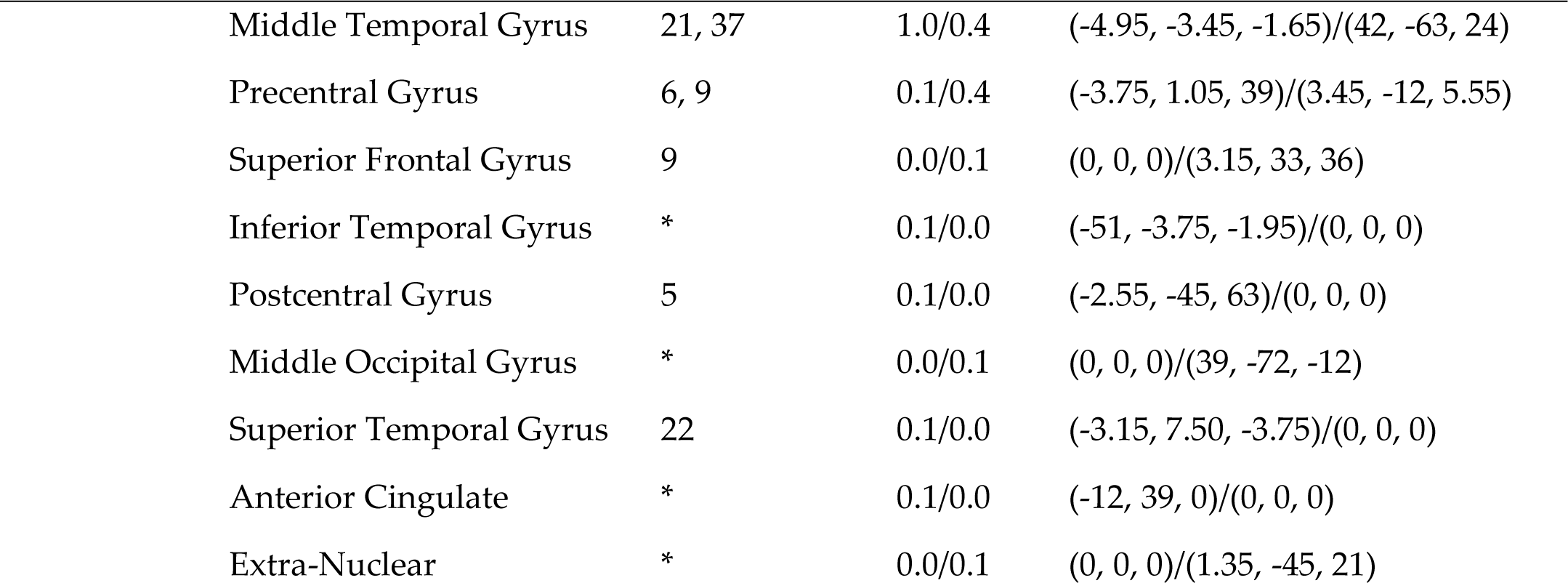
Talairach coordinates the brain circuit associated with narcissism.

## 4. Predictive model results

The supervised learning algorithm by mean of random forest regression results confirmed that the model was able to cover a quarter of variance with R^2^ of 0.247 (MSE= 1.098, RMSE=1.048; MAE=0.876). The validation error was of 0.835 MSE, and the out of bag (OOB) error of 0.771 MSE. The number of trees to reach optimal performance was 23 with two features per split. The mean decrease in purity confirmed the importance of IC-GM2 as a main predictor and main split in trees (see Figure 5).

**Figure 5.**
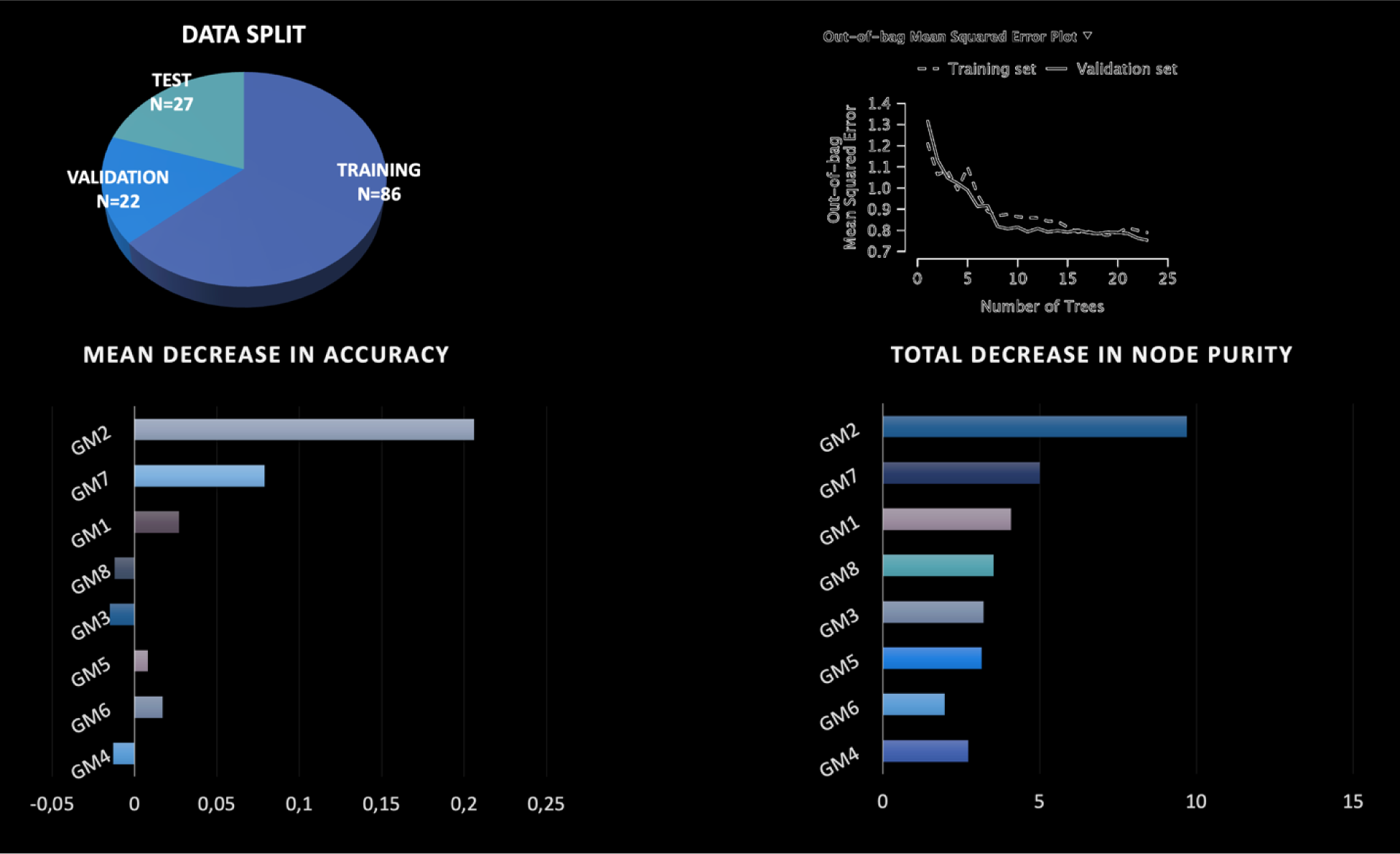
Random Forest regression results. The model confirmed the same circuit of stepwise regression was the best predictor of narcissistic personality traits.

## 5. Discussion

The primary goal of our study was to delineate the joint contributions of GM and WM to NPT (aim 1). Additionally, we aimed to scrutinize the specificity of our findings concerning the prediction of NPT, distinct from other related personality traits (aim 2). Lastly, we sought to assess the generalizability of our results by constructing a predictive model (aim 3). To achieve these objectives, we initially employed a data fusion unsupervised machine learning algorithm to analyze the structural MRI images of 135 healthy individuals. This approach facilitated the decomposition of the brain into independent networks characterized by the co-variation of GM and WM. Subsequently, we utilized stepwise regression and Random Forest regression to identify the GM-WM network associated with narcissistic traits and to extract a predictive model. The data fusion ML we employed detected a total of eight GM-WM brain networks, among which only IC-GM2/WM3 demonstrated a significant association with narcissistic personality traits. IC-GM2 encompassed regions such as the frontal-insular, temporal and parietal lobes, parieto-occipital areas, and limbic regions largely overlapping with the Default Mode Network. Although our data are structural and non-resting state, several previous studies have shown that resting state macro-networks were also present at a structural level when data are analyzed with ICA-based methods (Baggio et al., 2023; Grecucci et al., 2022; Meier et al., 2016; Vanasse et al., 2021). Meanwhile, IC-WM3 included portions of the fronto-parietal, thalamus, and cerebellum regions. Notably, this particular network did not predict any other personality trait as we examined and emerged as the most robust predictor within a Random Forest regression model. Additional details regarding these findings are expounded upon in the subsequent section.

The main areas encompassed within IC-GM2 comprise frontal, temporal, and parietal regions, including the superior temporal gyrus, the angular gyrus, the supramarginal gyrus, the middle temporal gyrus, and the inferior parietal lobe. These findings align with our earlier study, which also identified parietal and temporal areas, such as the angular gyrus, the rolandic operculum, and the Heschl’s gyrus, as relevant to narcissism (Jornkokgoud et al., 2023). These areas likely play a pivotal role in empathic and social cognitive processing linked to narcissism. Specifically, while the superior temporal gyrus and Heschl’s gyrus are primarily associated with auditory processing and language, they have also been implicated in social cognition (Bigler et al., 2007; Howard et al., 2000). Furthermore, the temporal and parietal lobes, including the superior temporal sulcus (STS) and angular gyrus (AG), demarcate the boundaries of the temporo–parietal junction (TPJ) regions (Schurz et al., 2017). This suggests the simultaneous engagement of social-cognitive and affective processes (Schurz et al., 2021). Narcissistic individuals, who employ socio-cognitive, emotional, intrapersonal, and interpersonal self-regulatory mechanisms to construct and uphold their disproportionately positive self-views, may manifest distinctive social interaction patterns (Eddy, 2023).

In this regard, Schmälzle et al. (2017) delved into the intricacies of brain connectivity during social interactions, aiming to map the structure of social networks. Their investigation unearthed shifts in connectivity patterns within brain regions associated with the mentalizing system, particularly as a coping mechanism for dealing with exclusion. Notably, individuals who exhibited heightened connectivity between the left and right temporoparietal junction (TPJ), two integral components of the mentalizing system, following instances of exclusion, tended to exhibit less intricate social network structures (Schmälzle et al., 2017). This aligns with the findings of Feng et al. (2021), who, in a comprehensive neuroimaging meta-analysis, underscored the role of the TPJ within the default mode network during diverse social interactions. This suggests that social cognitive processes form a vital foundation for human social relationships.

Furthermore, the current findings highlight the significance of the frontal region, the cingulate gyrus, and the insular gray matter to narcissism. These observations align with prior scientific research that has established a network involving the frontal and temporal lobes, including the insular cortex, in processes related to empathy (Iacoboni & Lenzi, 2002). Recent studies have shed light on the crucial roles played by the cingulate and insula in emotional processing and cognitive control, highlighting their activity and functional connectivity (Dupont et al., 2023). In particular, narcissism has been linked to functional deficits within the anterior insula and anterior cingulate cortex, which can result in challenges in both comprehending and sharing the emotions of others (Cascio et al., 2014; Jankowiak-Siuda & Zajkowski, 2013). Moreover, within the IC-GM2 network, other significant regions were detected in the parieto-occipital areas, including the middle occipital gyrus and the cuneus. Previous research in functional brain imaging has demonstrated the involvement of these areas in processing empathy during social interactions (Hamada et al., 2022). In fact, individuals with heightened levels of empathy tend to exhibit increased cortical activities within the parieto-occipital area (Hamada et al., 2022).

Of note, IC-GM2 largely overlaps with the DMN. The DMN is known to be activated during rest and is associated with anxiety, rumination, personality disorders and other psychopathologies (Vicentini et al., 2017; Zhou et al., 2020; Baggio et al., 2023; Langerbeck et al., 2023). Previous studies have confirmed abnormal functional connectivity within the DMN in psychological disorders (O’Neill et al., 2015; Wolf et al., 2011). The current results confirms the role of DMN for narcissistic personality as well. As for the white matter network, IC-WM3 predominantly encompassed regions within the cerebellum, suggesting a potential disruption in cerebellar connections. The cerebellar areas have gained significant attention in contemporary neuroscience due to their involvement in various cognitive and affective functions. For instance, social cognition has been closely linked to cerebellar functioning (Schmahmann, 2019), In a parallel manner to the processes associated with narcissism, there is evidence indicating that the propensity to seek novel experiences that elicit strong emotional reactions, often referred to as novelty-seeking, correlates with cerebellar white matter volumes (Picerni et al., 2013).

Deficits in the cerebellum have been closely associated with various psychopathological conditions (Kim et al., 2021), primarily due to their role in information integration, coordination, and monitoring (Hariri, 2019). Specifically, the cerebellum is a crucial component of the broader cerebello-thalamo-cortical circuit (CTCC) implicated in executive control functions, and its structural integrity may serve as a transdiagnostic marker for the risk of multiple psychopathologies (Hariri, 2019). Within this circuit, the thalamus plays a pivotal role in integrating stimuli originating from both the cerebellum and the cortex, as well as engaging in multisensory processing (Tyll et al., 2011). In studies examining narcissism, white matter irregularities in thalamic tracts, including the anterior thalamic radiation (Nenadić et al., 2021; Schmidt et al., 2023) and the posterior thalamic radiation (Lou et al., 2023), have been identified, substantiating our results.

Other regions inside the IC-WM3 were the precuneus, extending to the anterior cingulate, as well lateral temporal lobe areas, including middle and inferior temporal gyrus, white matter. All these regions have been shown to support the DMN (Teipel et al., 2010). Recent investigations have shed light on the involvement of parietal and frontal-limbic tracts in narcissism, possibly in relation to neuroticism, negative emotionality, and social subordination phenotypes (Nenadić et al., 2021; Schmidt et al., 2023). As emphasized in the introduction, our initial expectation was that the network of brain regions characterizing narcissistic personality traits would distinguish itself from those predominantly linked to other related yet distinct personality traits, such as histrionic (Cluster B), avoidant (Cluster C), and paranoid (Cluster A) traits.

Our findings align with this expectation, underscoring the presence of a specific brain network for narcissistic traits that is not shared with other personality profiles. Specifically, when exploring histrionic traits, which fall within the same Cluster B category as narcissistic traits, our research did not yield statistically significant outcomes. Histrionic and Narcissistic Personality Disorder (NPD) individuals are often characterized by their desire to be the center of attention, heightened emotional expression, and dramatic or unpredictable behaviors (American Psychiatric Association, 2013). Scientific investigations into the association between brain irregularities and histrionic traits remain limited. In a recent study, Langerbeck et al. (2023) employed supervised learning to construct a predictive model for borderline and histrionic traits using structural brain data. Interestingly, their findings suggested that borderline and histrionic traits do not share the same neural circuitry. These results, consistent with our study, indicate that personality traits within Cluster B are reliant on distinct brain networks.

In relation to avoidant personality traits, our study revealed that IC-GM1 and IC-GM5, but not IC-GM2, emerged as predictors within the model (see supplementary material). These two networks are likely associated with social inhibitions, feelings of inadequacy, and hypersensitivity to negative judgment, all of which are characteristic features often observed in individuals with avoidant personality disorder (APD) (American Psychiatric Association, 2013). Consequently, these traits may also manifest in individuals displaying avoidant personality traits. Although there are certain similarities between avoidant personality disorder and narcissistic personality disorder, such as insecure attachment patterns (Simonsen & Euler, 2019), and the avoidance of social situations in which they can be ashamed (Ronningstam, 2020), we found no overlap at a neural level, confirming different neural substrates in the two disorders. Moreover, our earlier findings demonstrated a significant relationship between insecure personality traits and the level of narcissism (Jornkokgoud et al., 2023), likely attributed to the connection between vulnerable narcissism and individuals with insecure tendencies (Kowalchyk et al., 2021).

Last but not least, for paranoid personality traits, the predictive factors that remained in the model included IC-GM3 and IC-GM5. Once again, the neural networks that predict paranoid personality traits were distinct from those associated with narcissistic traits, indicating the presence of unique neural mechanisms governing these two personality traits. Overall, the main results of the current study provide evidence supporting the existence of separate brain networks characterizing narcissistic, histrionic, avoidant, and paranoid personality traits, suggesting unique neural signatures for each trait.

Finally, we were able to extract a predictive model of NPT via Random Forest supervised ML. Results showed that the same network was also predictive of new cases via the cross-validation method. In support of these findings, Random Forest regression is considered a valuable forecasting technique that gains insight into its predictive performance through the framework’s analysis of individual predictor correlations and strengths. Thus, this technique is able to translate theoretical values into reality with the use of out-of-bag estimates (Breiman, 2001). It is characterized by its robustness against overfitting and its excellent suitability for predictive MRI data analysis (Sarica et al., 2017). In a similar vien, previous studies have also shown the possibility of predicting the diagnoses of borderline personality disorder cases from gray and white matter networks by using random forest classification (Grecucci, Dadomo, et al., 2023). Moreover, meaningful brain regions can be identified by using feature importances provided by random forest that shows diagnostic classification and investigating biomarkers from MRI features in other clinical diseases such as schizophrenia and bipolar disorder (Schwarz et al., 2019), Alzheimer’s disease (Song et al., 2021), neuromyelitis optica and multiple sclerosis (Eshaghi et al., 2016). Thus, it is safe to conclude that the current finding supports the use of the supervised machine learning, known as Random Forest regression, in which results for new observational cases accurately predicted from MRI data parameters, especially with gray and white matter.

## 5. Conclusions and limitations

This study represents the first attempt to employ unsupervised machine learning and correlation analysis of covarying GM-WM networks to test the hypothesis that distinct brain networks can predict individual variations in narcissistic personality traits. Our findings have successfully established a predictive model for narcissistic traits based on both gray and white matter features. Additionally, these results have furnished compelling evidence for the existence of a specific neural circuit associated with narcissism, setting it apart from other personality traits, whether within the same or different personality clusters, by using supervised machine learning such as Random Forest regression.

Despite the merits, this research does not come without limitations. Firstly, the findings are confined to the features of gray and white matter, with functional data not being considered at this time. Future investigations may benefit from incorporating functional imaging techniques to gain a more comprehensive understanding of the neural mechanisms underlying narcissism. Additionally, although the sample size in this study surpassed that of previous research, a further increase in sample size could be beneficial for brain-wide association analysis in subsequent studies (Marek et al., 2022). Another noteworthy issue pertains to the assessment of narcissistic traits. In our study, narcissism was gauged using the PSDI, which does not offer a clear distinction between vulnerable and grandiose subtypes. Future research should delve into this differentiation and explore potential variations between these two manifestations of narcissism.

## 6. Data Availability Statement

The dataset analyzed during the current study is available in the MPI-Leipzig_Mind-Brain-Body repository, https://openneuro.org/datasets/ds000221/versions/1.0.0 (accessed on 1 April 2022). The complete LEMON Data can be accessed via Gesellschaft für wissenschaftliche Datenverarbeitung mbH Göttingen (GWDG) https://www.gwdg.de/ (accessed on 1 April 2022). Raw and preprocessed data at this location is accessible through web browser https://ftp.gwdg.de/pub/misc/MPI-Leipzig_Mind-Brain-Body-LEMON/ (accessed on 1 December 2021) and a fast FTP connection (ftp://ftp.gwdg.de/pub/misc/MPI-Leipzig_Mind-Brain-Body-LEMON/ (accessed on 1 April 2022)).

