## Supplementary material for "Narcissus reflected: gray and white matter features joint contribution to the default mode network in predicting narcissistic personality traits": Table 1

**Table 1. Correlation coefficients between the linked components**

| IC-GM | IC-WM | *r* | t-value | p-value |
|---|---|---|---|---|
| 8 | 3 | 0.524 | 7.100 | <0.001 |
| 1 | 3 | 0.429 | 5.472 | <0.001 |
| 5 | 5 | 0.345 | 4.242 | <0.001 |
| 4 | 3 | 0.338 | 4.139 | <0.001 |
| 3 | 5 | -0.303 | -3.672 | <0.001 |
| 7 | 7 | 0.299 | 3.619 | <0.001 |
| 6 | 8 | 0.294 | 3.546 | <0.001 |
| 2 | 3 | 0.247 | 2.936 | <0.001 |
