## Supplementary material for "Narcissus reflected: gray and white matter features joint contribution to the default mode network in predicting narcissistic personality traits": Table 2

**Table 2. Talairach coordinates the brain circuit associated with narcissism.**

| Component | Area | Brodmann Area | volume (cc) | MNI (x, y, z) |
|---|---|---|---|---|
| IC-GM 2 | | | | |
| Positive | | | | |
| | Superior Temporal Gyrus | 13, 22, 39, 41 | 4.3/4.0 | (-42, -55, 28)/(43, -56, 26) |
| | Angular Gyrus | 39 | 1.0/1.3 | (-43, -58, 31)/(43, -55, 32) |
| | Supramarginal Gyrus | 40 | 3.6/2.8 | (-48, -53, 23)/(40, -53, 26) |
| | Middle Temporal Gyrus | 21, 22, 39 | 5.5/1.4 | (-43, -56, 22)/(40, -56, 24) |
| | Inferior Parietal Lobule | 2, 39, 40 | 5.9/7.2 | (-45, -48, 23)/(36, -31, 42) |
| | Sub-Gyral | * | 2.6/2.9 | (-40, -48, 25)/ (39, -31, 44) |
| | Postcentral Gyrus | 1, 2, 3, 5, 7, 40 | 6.5/7.0 | (-49, -23, 36)/(48, -21, 40) |
| | Precentral Gyrus | 3, 4 | 0.6/1.7 | (-59, -13, 30)/(50, -19, 37) |
| | Cingulate Gyrus | 23, 24, 31 | 2.4/1.8 | (-4, -40, 38)/(3, -39, 38) |
| | Precuneus | 7, 19, 31, 39 | 5.1/4.6 | (-1, -45, 56)/(4, -47, 56) |
| | Paracentral Lobule | 4, 5 | 1.0/0.6 | (-3, -45, 60)/(3, -44, 62) |
| | Superior Parietal Lobule | 5, 7 | 1.7/0.8 | (-30, -65, 49)/(33, -68, 43) |
| | Middle Occipital Gyrus | * | 0.6/0.0 | (-45, -63, -4)/(0, 0, 0) |
| | Parahippocampal Gyrus | 19, 36, 37 | 0.7/0.5 | (-25, -47, -5)/(28, -45, -5) |
| | Lingual Gyrus | 18 | 0.3/0.0 | (-6, -67, 2)/(0, 0, 0) |
| | Insula | 13 | 0.0/0.1 | (0, 0, 0)/(46, -41, 20) |
| | Inferior Temporal Gyrus | 37 | 0.2/0.0 | (-43, -68, 1)/(0, 0, 0) |
| | Fusiform Gyrus | 19, 37 | 0.1/0.1 | (-25, -49, -9)/(27, -49, -9) |
| | Declive | * | 0.1/0.2 | (-27, -52, -11)/(28, -57, -14) |
| | Orbital Gyrus | * | 0.0/0.1 | (0, 0, 0)/(9, 47, -21) |

| Component | Area | Brodmann Area | volume (cc) | MNI (x, y, z) |
|---|---|---|---|---|

| Component | Area | Brodmann Area | volume (cc) | MNI (x, y, z) |
|---|---|---|---|---|
| IC-GM 2 | | | | |
| Negative | | | | |
| | Posterior Cingulate | 30, 31 | 1.2/1.0 | (-21, -64, 9)/(22, -61, 10) |
| | Cuneus | 17, 18, 23, 30 | 0.8/0.3 | (-18, -67, 9)/(21, -68, 10) |
| | Extra-Nuclear | * | 0.4/0.6 | (-24, -56, 8)/(24, -56, 8) |
| | Declive | * | 0.3/1.2 | (-12, -84, -22)/(10, -83, -21) |
| | Lateral Ventricle | * | 0.1/0.3 | (-25, -53, 11)/(27, -53, 8) |
| | Uvula | * | 0.8/0.6 | (-13, -84, -26)/(13, -83, -24) |
| | Pyramis | * | 0.8/0.5 | (-10, -82, -29)/(10, -80, -24) |
| | Parahippocampal Gyrus | 30 | 0.0/0.2 | (0, 0, 0)/(24, -54, 5) |
| | Sub-Gyral | * | 0.0/0.3 | (0, 0, 0)/(33, -49, 40) |
| | Middle Frontal Gyrus | 6 | 0.3/0.0 | (-16, -10, 61)/(0, 0, 0) |
| | Lingual Gyrus | * | 0.1/0.0 | (-21, -61, 4)/(0, 0, 0) |
| | Tuber | * | 0.1/0.0 | (-21, -83, -30)/(0, 0, 0) |
| | Medial Frontal Gyrus | * | 0.1/0.0 | (-13, -10, 64)/(0, 0, 0) |
| | Inferior Semi-Lunar Lobule | * | 0.1/0.0 | (-16, -79, -35)/(0, 0, 0) |
| | Superior Frontal Gyrus | 6 | 0.1/0.1 | (-16, -7, 64)/(24, 10, 51) |
| IC-WM 3 | | | | |
| Positive | | | | |
| | Cerebellar Tonsil | * | 4.9/4.5 | (-25, -54, -31)/(21, -47, -32) |
| | Culmen | * | 3.0/3.6 | (-21, -55, -25)/(24, -51, -26) |
| | Pyramis | * | 1.4/1.5 | (-16, -61, -27)/(21, -60, -27) |
| | Nodule | * | 0.5/0.6 | (-13, -51, -28)/(13, -57, -29) |
| | Tuber | * | 0.3/0.6 | (-31, -60, -27)/(30, -58, -30) |
| | Fastigium | * | 0.4/0.4 | (-10, -61, -22)/(10, -61, -22) |
| | Declive | * | 0.6/0.8 | (-13, -64, -21)/(13, -64, -22) |

| Component | Area | Brodmann Area | volume (cc) | MNI (x, y, z) |
|---|---|---|---|---|
| | Uvula | * | 0.8/0.5 | (-13, -64, -31)/(12, -61, -30) |
| | Extra-Nuclear | * | 1.8/1.2 | (-12, -9, -5)/(18, -15, -3) |
| | Lentiform Nucleus | * | 0.7/1.1 | (-15, -7, -2)/(18, -10, -1) |
| | Inferior Semi-Lunar Lobule | * | 0.3/0.3 | (-24, -64, -35)/(22, -63, -35) |
| | Thalamus | * | 0.3/0.1 | (-15, -18, 0)/(16, -17, 1) |
| | Cerebellar Lingual | * | 0.1/0.0 | (-6, -46, -18)/(0, 0, 0) |
| | Inferior Parietal Lobule | 40 | 0.1/0.0 | (-40, -49, 38)/(0, 0, 0) |
| | Fourth Ventricle | * | 0.1/0.1 | (-3, -52, -23)/(1, -40, -19) |
| | Supramarginal Gyrus | 40 | 0.1/0.0 | (-39, -49, 34)/(0, 0, 0) |
| | Sub-Gyral | * | 0.1/0.0 | (-34, -32, 7)/(0, 0, 0) |

| Component | Area | Brodmann Area | volume (cc) | MNI (x, y, z) |
|---|---|---|---|---|
| IC-WM 3 Negative | | | | |
| | Sub-Gyral | * | 0.1/0.0 | (-30, -42, 46)/(0, 0, 0) |
| | Middle Frontal Gyrus | 9 | 0.1/0.4 | (-4.05, 1.05, 42)/(2.85, 30, 36) |
| | Paracentral Lobule | 5 | 0.2/0.1 | (-1.95, -45, 51)/(15, -36, 57) |
| | Superior Parietal Lobule | 7 | 0.1/0.4 | (-12, -6.45, 54)/(2.55, -57, 4.65) |
| | Inferior Parietal Lobule | 40 | 0.6/0.1 | (-33, -45, 5.25)/(3.15, -57, 4.65) |
| | Precuneus | 7 | 1.9/1.2 | (-12, -6.45, 48)/(2.85, -57, 4.95) |
| | Uncus | 20 | 0.6/0.1 | (-3.45, -6, -3.75)/(30, 4.50, -36) |
| | Middle Temporal Gyrus | 21, 37 | 1.0/0.4 | (-4.95, -3.45, -1.65)/(42, -63, 24) |
| | Precentral Gyrus | 6, 9 | 0.1/0.4 | (-3.75, 1.05, 39)/(3.45, -12, 5.55) |
| | Superior Frontal Gyrus | 9 | 0.0/0.1 | (0, 0, 0)/(3.15, 33, 36) |
| | Inferior Temporal Gyrus | * | 0.1/0.0 | (-51, -3.75, -1.95)/(0, 0, 0) |
| | Postcentral Gyrus | 5 | 0.1/0.0 | (-2.55, -45, 63)/(0, 0, 0) |

| Component | Area | Brodmann Area | volume (cc) | MNI (x, y, z) |
|---|---|---|---|---|
| | Middle Occipital Gyrus | * | 0.0/0.1 | (0, 0, 0)/(39, -72, -12) |
| | Superior Temporal Gyrus | 22 | 0.1/0.0 | (-3.15, 7.50, -3.75)/(0, 0, 0) |
| | Anterior Cingulate | * | 0.1/0.0 | (-12, 39, 0)/(0, 0, 0) |
| | Extra-Nuclear | * | 0.0/0.1 | (0, 0, 0)/(1.35, -45, 21) |
