## Supplementary material for "Narcissus reflected: gray and white matter features joint contribution to the default mode network in predicting narcissistic personality traits": IC-WM3

### Supplementary figures and tables

**Figure 6.**

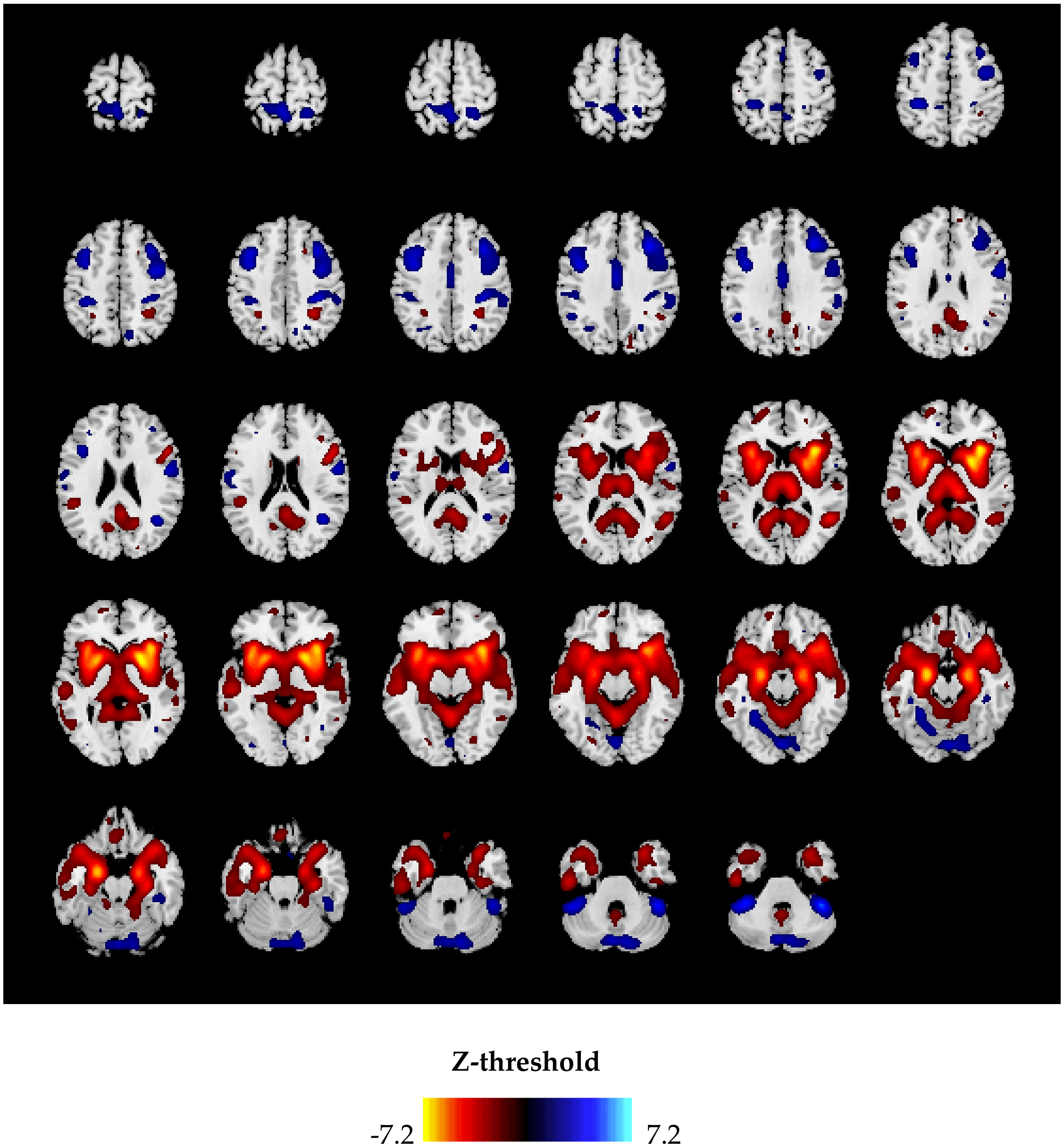


Figure 6. Gray matter independent component 1 (IC-GM1). Warm hues show positive value areas, whereas cool colors represent negative regions.

**Figure 7.**

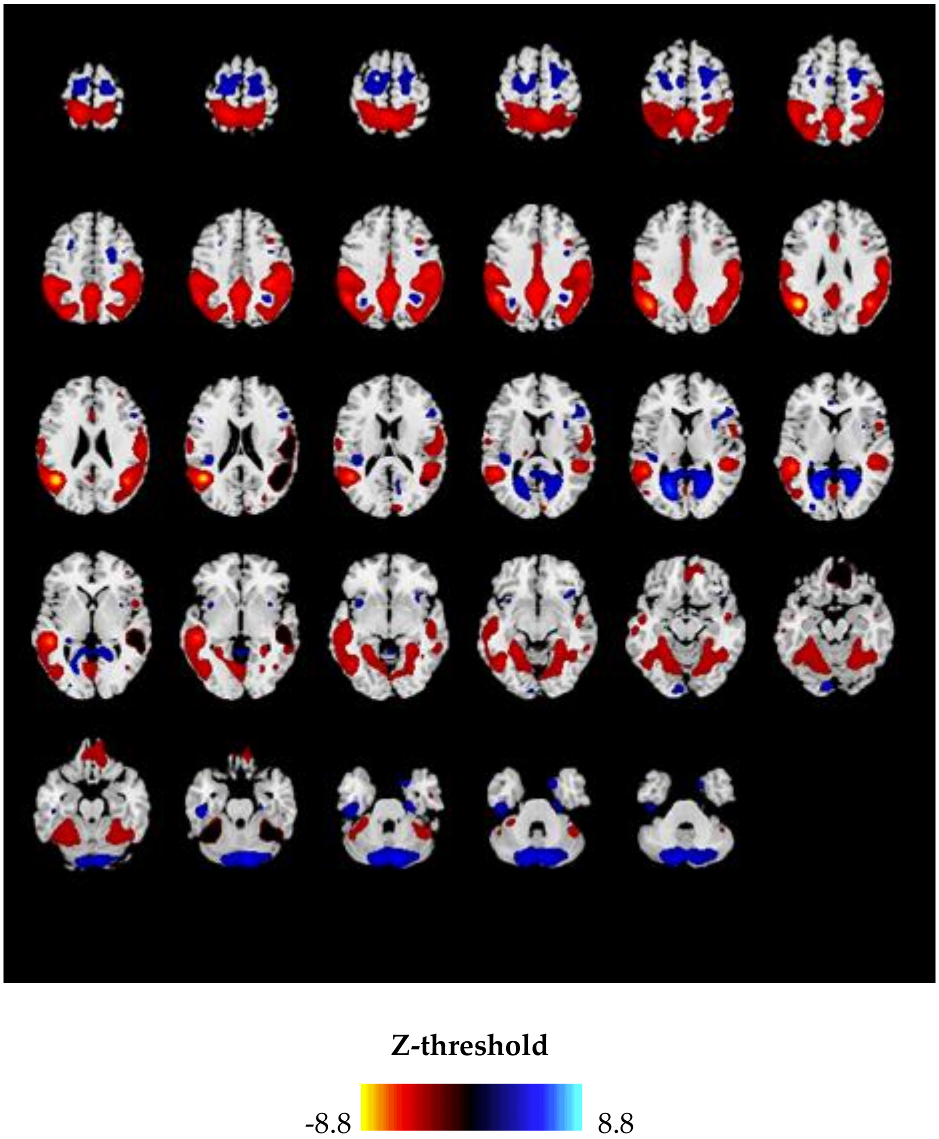


Figure 7. Gray matter independent component 2 (IC-GM2). Warm hues show positive value areas, whereas cool colors represent negative regions.

**Figure 8.**

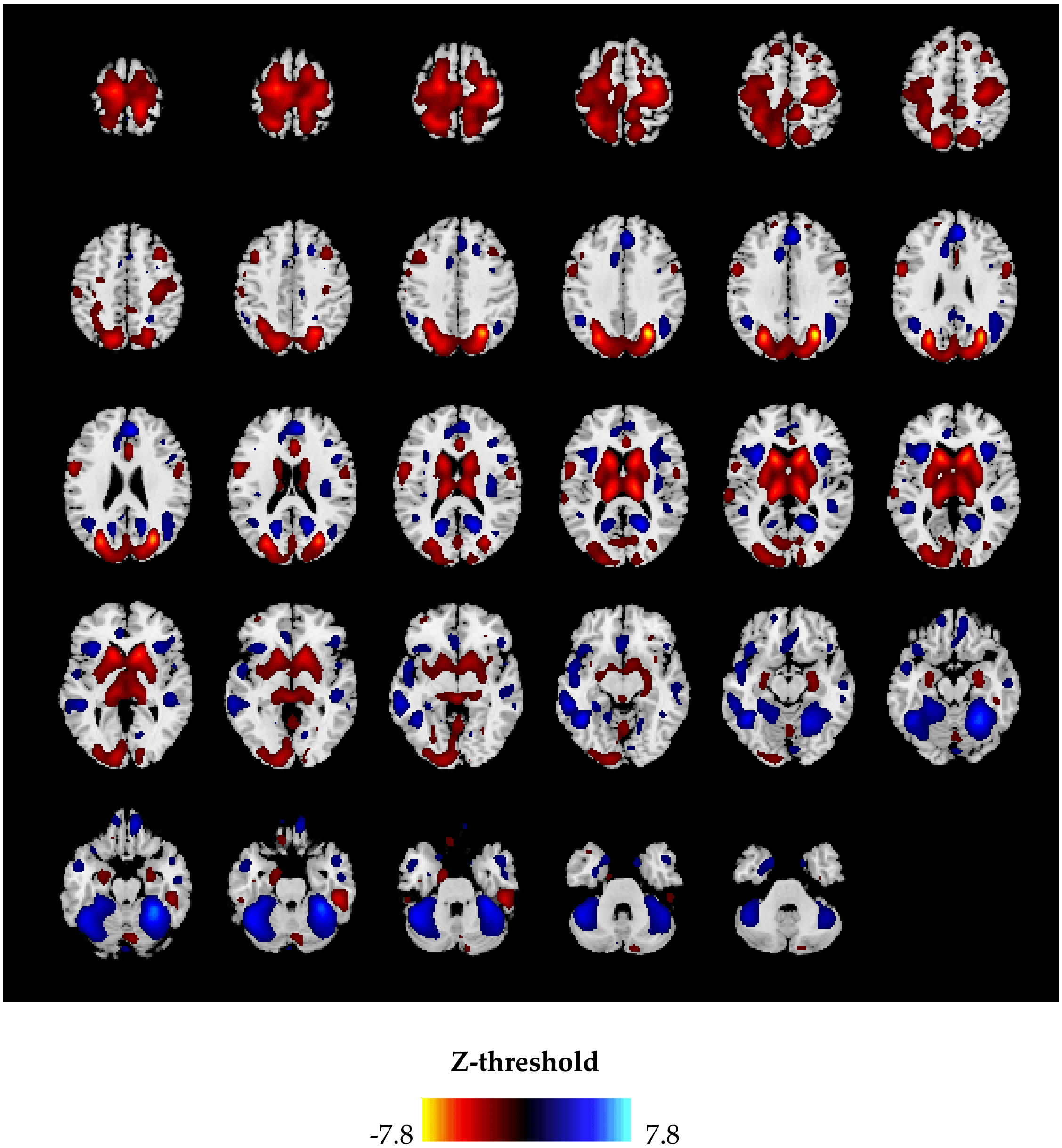


Figure 8. Gray matter independent component 3 (IC-GM3). Warm hues show positive value areas, whereas cool colors represent negative regions.

**Figure 9.**

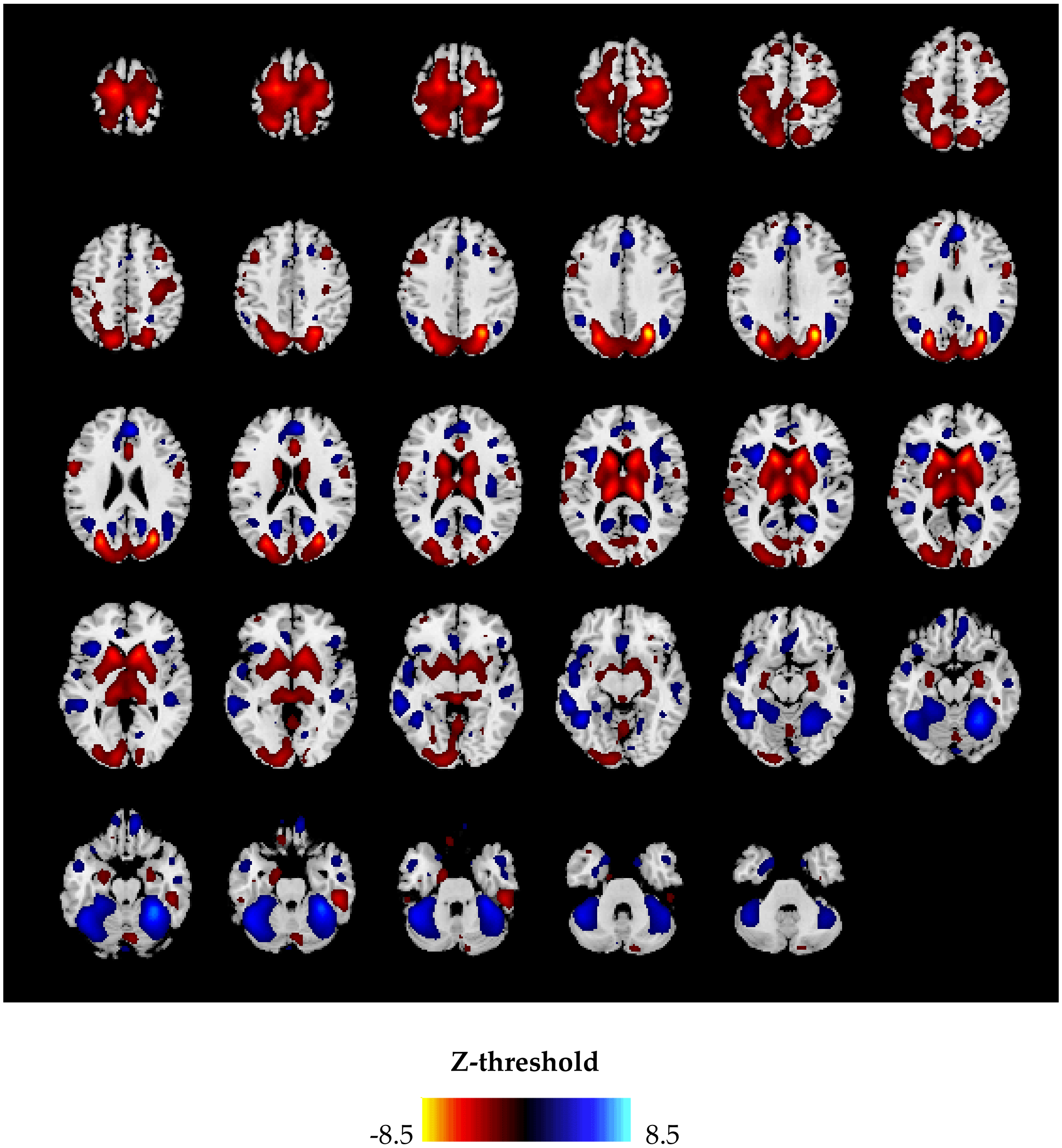


Figure 9. Gray matter independent component 5 (IC-GM5). Warm hues show positive value areas, whereas cool colors represent negative regions.

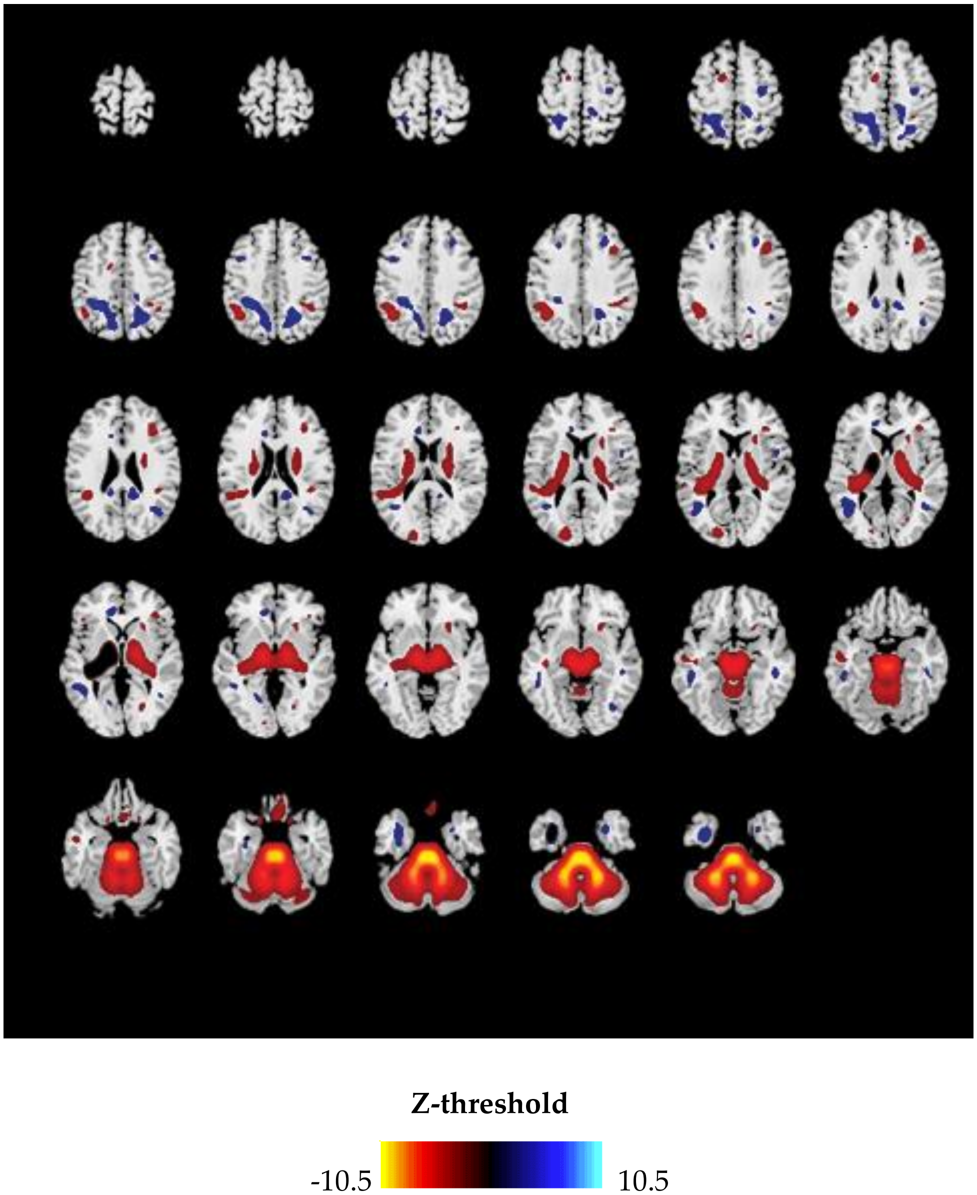


Figure 9. White matter independent component 3 (IC-WM3). Warm hues show positive value areas, whereas cool colors represent negative regions.

**Table 3. Talairach coordinates the brain circuit associated with other personality traits**

| Component | Area | Brodmann Area | volume (cc) | MNI (x, y, z) |
|---|---|---|---|---|
| IC-GM 1 | | | | |
| Positive | | | | |
| | Insula | 13 | 3.1/4.4 | (-36, 1.35, -6)/(36, 1.95, 6) |
| | Lentiform Nucleus | * | 3.6/4.1 | (-2.25, 9, -1.50)/(2.55, 9, 0) |
| | Parahippocampal Gyrus | 28, 30, 34, 35, 36 | 6.7/5.1 | (-2.25, -7.50, -21)/(2.25, -9, -1.95) |
| | Inferior Frontal Gyrus | 13, 44, 45, 47 | 2.2/5.6 | (-3.45, 1.65, -3)/(36, 18, -6) |
| | Extra-Nuclear | 13, 47 | 6.2/6.0 | (-33, 1.65, -7.50)/(2.85, 1.05, 3) |
| | Uncus | 28, 34 | 1.5/0.7 | (-2.25, -4.50, -24)/(2.25, -6, -24) |
| | Claustrum | * | 0.6/0.8 | (-33, 7.50, -1.50)/(3.15, 12, 6) |
| | Lateral Ventricle | * | 0.5/0.2 | (-2.55, -7.50, -24)/(27, -6, -2.55) |
| | Superior Temporal Gyrus | 22, 38, 39 | 3.1/3.7 | (-42, 1.65, -2.25)/(4.35, 18, -18) |
| | Precentral Gyrus | 44 | 0.0/0.1 | (0, 0, 0)/(4.35, 1.95, 9) |
| | Caudate | * | 0.4/0.9 | (-1.35, 1.65, -1.50)/(1.65, 15, 1.50) |
| | Thalamus | * | 1.5/3.0 | (-1.05, -27, 6)/(12, -27, 4.50) |
| | * | * | 0.4/0.6 | (-1.95, -7.50, -12)/(4.35, 24, -18) |
| | Sub-Gyral | * | 0.8/0.9 | (-33, 2.55, 0)/(3.15, 27, 1.50) |
| | Culmen | * | 1.2/0.7 | (0, -5.25, -9)/(3, -54, -12) |
| | Subcallosal Gyrus | 34 | 0.3/0.8 | (-2.25, 4.50, -15)/(1.65, 1.05, -1.35) |
| | Anterior Cingulate | 25 | 0.4/0.4 | (-1.50, 4.50, -6)/(6, 9, -7.50) |
| | Posterior Cingulate | 23, 29, 30 | 0.6/0.6 | (-1.65, -57, 6)/(9, -5.25, 9) |
| | Middle Temporal Gyrus | 21 | 0.2/0.5 | (-57, -2.25, -4.50)/(5.25, -5.25, 6) |
| | Medial Frontal Gyrus | 25 | 0.0/0.1 | (0, 0, 0)/(15, 12, -18) |

| Component | Area | Brodmann Area | volume (cc) | MNI (x, y, z) |
| --- | --- | --- | --- | --- |
| | Third Ventricle | * | 0.1/0.1 | (-1.50, -1.65, 1.50)/(1.50, -9, 1.50) |
| | Lingual Gyrus | 18 | 0.1/0.0 | (-1.65, -54, 3)/(0, 0, 0) |
| IC-GM 1 Negative | | | | |
| | Cerebellar Tonsil | * | 3.1/2.6 | (-3.75, -45, -4.95)/(42, -45, -4.65) |
| | Culmen | * | 0.3/0.5 | (-4.05, -42, -3.75)/(45, -45, -3.75) |
| | Middle Frontal Gyrus | 9 | 0.1/0.6 | (-36, 1.65, 4.35)/(36, 2.55, 3.45) |
| | Inferior Semi-Lunar Lobule | * | 0.3/0.0 | (-15, -6.75, -5.25)/(0, 0, 0) |
| | Inferior Frontal Gyrus | * | 0.1/0.0 | (-4.05, 6, 36)/(0, 0, 0) |
| | Superior Temporal Gyrus | 39 | 0.0/0.1 | (0, 0, 0)/(4.35, -5.55, 21) |
| | Postcentral Gyrus | 2 | 0.0/0.1 | (0, 0, 0)/(36, -3.15, 42) |
| | Precentral Gyrus | * | 0.1/0.0 | (-39, 6, 4.05)/(0, 0, 0) |

| Component | Area | Brodmann Area | volume (cc) | MNI (x, y, z) |
| --- | --- | --- | --- | --- |
| IC-GM 3 Positive | | | | |
| | Precuneus | 7, 19, 31 | 4.3/3.6 | (-2.55, -69, 33)/(30, -6.75, 33) |
| | Sub-Gyral | 6 | 3.2/3.3 | (-27, -72, 30)/(30, -7.05, 30) |
| | Caudate | * | 2.4/2.4 | (-1.35, 15, 6)/(1.35, 15, 6) |
| | Thalamus | * | 3.3/3.6 | (-1.35, -2.25, 9)/(1.35, -1.95, 1.05) |
| | Precentral Gyrus | 4, 6 | 1.9/5.6 | (-1.05, -1.95, 72)/(36, -12, 6.15) |
| | Superior Frontal Gyrus | 6 | 0.9/0.1 | (-1.65, -15, 6.75)/(30, -12, 6.75) |

| Component | Area | Brodmann Area | volume (cc) | MNI (x, y, z) |
|---|---|---|---|---|
| | Middle Frontal Gyrus | 6 | 1.3/1.2 | (-18, -12, 6.45)/(33, -9, 6.15) |
| | Superior Parietal Lobule | 7 | 1.5/0.9 | (-30, -4.95, 6.15)/(2.85, -6.75, 4.35) |
| | Extra-Nuclear | * | 1.3/1.4 | (-1.65, 12, 6)/(18, 21, 4.50) |
| | Medial Frontal Gyrus | 6 | 1.3/0.0 | (-1.05, -1.35, 7.05)/(0, 0, 0) |
| | * | * | 0.0/0.0 | (0, 0, 0)/(0, 0, 0) |
| | Cuneus | 7, 17, 18, 19 | 1.3/1.3 | (-24, -84, 21)/(27, -7.95, 2.85) |
| | Superior Occipital Gyrus | * | 0.1/0.0 | (-30, -81, 24)/(0, 0, 0) |
| | Postcentral Gyrus | 3, 5, 7 | 1.3/1.2 | (-21, -54, 66)/(18, -42, 69) |
| | Lentiform Nucleus | * | 1.2/2.0 | (-1.95, 12, 9)/(1.95, 1.65, 0) |
| | Inferior Parietal Lobule | 40 | 0.6/0.0 | (-33, -4.95, 5.85)/(0, 0, 0) |
| | Angular Gyrus | * | 0.1/0.1 | (-3.15, -60, 36)/(33, -6.15, 36) |
| | Fusiform Gyrus | 20 | 0.0/0.3 | (0, 0, 0)/(5.25, -3.45, -2.85) |
| | Middle Occipital Gyrus | 19 | 0.4/0.0 | (-30, -81, 18)/(0, 0, 0) |
| | Inferior Frontal Gyrus | 9, 44 | 0.2/0.1 | (-57, 4.50, 27)/(5.85, 7.50, 30) |
| | Paracentral Lobule | 5, 6 | 0.1/0.3 | (-6, -3.45, 69)/(1.05, -42, 57) |
| | Lateral Ventricle | * | 0.0/0.1 | (0, 0, 0)/(1.65, 24, 7.50) |
| | Anterior Cingulate | 24 | 0.0/0.2 | (0, 0, 0)/(4.50, 2.85, 1.95) |
| | Middle Temporal Gyrus | * | 0.1/0.3 | (-30, -75, 18)/(33, -7.05, 1.95) |
| | Cerebellar Tonsil | * | 0.1/0.0 | (-27, -4.35, -5.25)/(0, 0, 0) |
| | Lingual Gyrus | 17 | 0.3/0.0 | (-7.50, -96, -3)/(0, 0, 0) |
| IC-GM 3 Negative | | | | |
| | Culmen | * | 3.1/4.4 | (-4.05, -5.25, -2.85)/(3.15, -48, -24) |
| | Declive | * | 1.5/2.6 | (-4.35, -5.55, -2.85)/(27, -5.55, -21) |

| Component | Area | Brodmann Area | volume (cc) | MNI (x, y, z) |
|---|---|---|---|---|
| | Fusiform Gyrus | 19, 20, 37 | 0.6/0.8 | (-45, -54, -18)/(36, -45, -24) |
| | * | * | 0.0/0.1 | (0, 0, 0)/(3.75, -51, -21) |
| | Medial Frontal Gyrus | 9 | 0.0/1.7 | (0, 0, 0)/(6, 45, 2.55) |
| | Sub-Gyral | * | 1.0/0.3 | (-45, -54, -1.35)/(1.95, -60, 1.95) |
| | Posterior Cingulate | 30 | 0.1/0.8 | (-1.65, -60, 15)/(21, -5.85, 1.05) |
| | Tuber | * | 1.0/0.3 | (-4.35, -5.25, -3.15)/(4.35, -5.55, -3.15) |
| | Extra-Nuclear | * | 0.1/0.6 | (-36, 24, 0)/(24, -5.85, 7.50) |
| | Insula | 13, 45 | 0.6/0.1 | (-33, 21, 6)/(36, 2.25, 6) |
| | Middle Temporal Gyrus | 21 | 0.6/0.0 | (-5.85, -39, -12)/(0, 0, 0) |
| | Inferior Temporal Gyrus | * | 0.1/0.0 | (-4.95, -54, -1.65)/(0, 0, 0) |
| | Inferior Frontal Gyrus | 13, 45, 47 | 0.3/0.1 | (-36, 21, 9)/(36, 2.55, 3) |
| | Rectal Gyrus | 11 | 0.0/0.1 | (0, 0, 0)/(1.05, 5.25, -24) |
| | Precuneus | * | 0.1/0.1 | (-1.65, -66, 21)/(1.65, -63, 21) |
| | Supramarginal Gyrus | * | 0.1/0.0 | (-45, -5.25, 33)/(0, 0, 0) |

| Component | Area | Brodmann Area | volume (cc) | MNI (x, y, z) |
|---|---|---|---|---|
| IC-GM 5 Positive | | | | |
| | Uncus | 20, 28, 34, 36, 38 | 3.6/1.9 | (-18, 0, -3.45)/(18, -1.50, -33) |
| | * | * | 0.1/0.2 | (-3, -8.85, -21)/(4.50, -90, -18) |

| Component | Area | Brodmann Area | volume (cc) | MNI (x, y, z) |
|---|---|---|---|---|
| | Lingual Gyrus | 17, 18 | 0.2/3.8 | (0, -90, -15)/(6, -90, -1.35) |
| | Parahippocampal Gyrus | 28, 34, 35 | 0.8/0.6 | (-21, -6, -3.15)/(21, -7.50, -30) |
| | Precuneus | 7, 31 | 3.1/2.0 | (0, -60, 3.75)/(3, -60, 4.05) |
| | Inferior Occipital Gyrus | 17, 18, 19 | 0.0/1.9 | (0, 0, 0)/(12, -93, -1.65) |
| | Middle Occipital Gyrus | 18, 19, 37 | 0.0/1.7 | (0, 0, 0)/(4.35, -75, -1.65) |
| | Paracentral Lobule | 4, 5, 6 | 0.8/1.8 | (0, -3.45, 6.45)/(3, -3.45, 6.75) |
| | Posterior Cingulate | 23, 29, 30, 31 | 1.2/0.6 | (-1.50, -5.25, 1.95)/(3, -5.25, 18) |
| | Cingulate Gyrus | 31 | 0.8/0.3 | (0, -57, 2.55)/(3, -54, 27) |
| | Medial Frontal Gyrus | 6 | 1.2/1.5 | (-1.50, -1.95, 72)/(3, -2.85, 6.15) |
| | Fusiform Gyrus | 19, 37 | 0.2/1.8 | (-4.35, -7.35, -18)/(45, -72, -1.95) |
| | Cuneus | 17, 18 | 0.0/2.0 | (0, 0, 0)/(6, -96, -1.50) |
| | Sub-Gyral | * | 0.3/0.1 | (-27, 3, -4.05)/(45, -69, -6) |
| | Lateral Ventricle | * | 0.1/0.1 | (-21, -6, -2.55)/(24, -7.50, -27) |
| | Cerebellar Tonsil | * | 0.0/0.5 | (0, 0, 0)/(3.15, -4.95, -5.25) |
| | Superior Temporal Gyrus | 38 | 0.1/0.0 | (-21, 9, -4.05)/(0, 0, 0) |
| | Declive | * | 0.0/0.2 | (0, 0, 0)/(4.35, -75, -2.25) |
| | Superior Parietal Lobule | 7 | 0.0/0.2 | (0, 0, 0)/(2.25, -66, 57) |
| | Inferior Temporal Gyrus | * | 0.0/0.1 | (0, 0, 0)/(45, -72, -3) |
| | Superior Frontal Gyrus | * | 0.1/0.1 | (-3, -7.50, 72)/(3, -1.05, 69) |
| | Extra-Nuclear | * | 0.1/0.1 | (-4.50, -45, 1.65)/(3, -4.35, 1.35) |
| | Middle Temporal Gyrus | 39 | 0.0/0.1 | (0, 0, 0)/(42, -7.65, 27) |
| | Precentral Gyrus | * | 0.0/0.1 | (0, 0, 0)/(1.05, -2.85, 6.75) |

| Component | Area | Brodmann Area | volume (cc) | MNI (x, y, z) |
|---|---|---|---|---|
| | Insula | * | 0.0/0.1 | (0, 0, 0)/(42, -1.95, 1.95) |
| IC-GM 5 Negative | | | | |
| | Superior Frontal Gyrus | 9, 10, 11 | 3.6/3.8 | (-24, 48, -15)/(30, 4.35, -15) |
| | Middle Frontal Gyrus | 6, 9, 11, 47 | 5.1/3.8 | (-27, 39, -15)/(33, 4.05, -15) |
| | * | * | 0.6/0.7 | (-36, 42, -15)/(36, 4.35, -15) |
| | Declive | * | 0.0/3.2 | (0, 0, 0)/(1.35, -81, -30) |
| | Inferior Frontal Gyrus | 11, 47 | 3.2/2.7 | (-2.55, 30, -1.35)/(2.85, 33, -15) |
| | Pyramis | * | 0.0/1.3 | (0, 0, 0)/(1.05, -81, -33) |
| | Uvula | * | 0.0/1.0 | (0, 0, 0)/(1.65, -81, -33) |
| | Sub-Gyral | * | 2.6/2.0 | (-3.45, -4.65, 4.35)/(2.25, 3.15, -15) |
| | Inferior Parietal Lobule | 7, 40 | 3.3/0.5 | (-36, -51, 45)/(3.75, -33, 4.35) |
| | Inferior Temporal Gyrus | 20 | 0.3/0.0 | (-4.65, -1.95, -3.45)/(0, 0, 0) |
| | Fusiform Gyrus | 20 | 0.4/0.0 | (-45, -2.25, -3.15)/(0, 0, 0) |
| | Postcentral Gyrus | 2, 3, 40 | 0.0/0.6 | (0, 0, 0)/(4.05, -30, 4.35) |
| | Medial Frontal Gyrus | * | 0.3/0.1 | (-12, 60, -18)/(1.05, 57, -18) |
| | Orbital Gyrus | 11, 47 | 0.2/0.2 | (-15, 2.25, -27)/(1.05, 54, -21) |
| | Middle Occipital Gyrus | * | 0.5/0.0 | (-36, -81, 12)/(0, 0, 0) |
| | Superior Parietal Lobule | 7 | 0.1/0.0 | (-30, -5.85, 45)/(0, 0, 0) |
| | Precuneus | * | 0.1/0.1 | (-2.85, -6.15, 42)/(1.65, -6.45, 36) |
| | Rectal Gyrus | * | 0.0/0.1 | (0, 0, 0)/(6, 51, -24) |
| | Precentral Gyrus | 44 | 0.2/0.0 | (-5.25, 1.05, 9)/(0, 0, 0) |
| | Supramarginal Gyrus | * | 0.1/0.1 | (-33, -5.55, 3.75)/(36, -54, 3.75) |
| | Cerebellar Tonsil | * | 0.1/0.0 | (-36, -6.45, -4.05)/(0, 0, 0) |
| | Culmen | * | 0.1/0.0 | (-3.45, -39, -3.15)/(0, 0, 0) |

| Component | Area | Brodmann Area | volume (cc) | MNI (x, y, z) |
|---|---|---|---|---|
| | Superior Temporal Gyrus | 39 | 0.0/0.1 | (0, 0, 0)/(4.65, -5.55, 1.05) |
| | Cuneus | 19 | 0.0/0.1 | (0, 0, 0)/(1.65, -84, 36) |
| | Middle Temporal Gyrus | * | 0.1/0.0 | (-3.75, -81, 1.65)/(0, 0, 0) |
| | Tuber | * | 0.1/0.0 | (-3.45, -69, -39)/(0, 0, 0) |
| | Parahippocampal Gyrus | 36 | 0.1/0.0 | (-4.05, -2.25, -2.25)/(0, 0, 0) |
